# Ultra-long-acting refillable nanofluidic implant confers full protection against SHIV infection in non-human primates

**DOI:** 10.1101/2022.12.15.520646

**Authors:** Fernanda P. Pons-Faudoa, Nicola Di Trani, Simone Capuani, Jocelyn Nikita Campa-Carranza, Bharti Nehete, Suman Sharma, Kathryn A. Shelton, Lane R. Bushman, Farah Abdelmawla, Martin Williams, Laura Roon, David Nerguizian, Corrine Ying Xuan Chua, Michael M. Ittmann, Joan E. Nichols, Jason T. Kimata, Peter L. Anderson, Pramod N. Nehete, Roberto C. Arduino, Alessandro Grattoni

## Abstract

The impact of pre-exposure prophylaxis (PrEP) on slowing the global human immunodeficiency virus (HIV) epidemic hinges on effective drugs and delivery platforms. Oral regimens have represented the pillar of HIV PrEP for years. However, variable adherence has spurred development of long-acting delivery systems, which also aim at increasing PrEP access, uptake and persistence. Here we present an ultra-long-acting and transcutaneously refillable subcutaneous nanofluidic implant for constant and sustained release of islatravir (ISL), a nucleoside reverse transcriptase translocation inhibitor, for HIV PrEP. In rhesus macaques, the ISL-eluting implants (nISL) achieved constant plasma ISL levels (median 3.14 nM) and peripheral blood mononuclear cells (PBMCs) ISL-triphosphate levels (ISL-TP) (median 0.16 pmol/10^6^ cells) for over 20 months uninterrupted. These drug concentrations are above the established PrEP protection threshold. In two non-blinded, placebo-controlled studies with repeated low-dose rectal and vaginal SHIV_SF162P3_ challenges in male and female rhesus macaques, respectively, nISL implants conferred 100% protection against infection (*p*=0.0005 and 0.0009, respectively between nISL and placebo control groups). The nISL implants were well tolerated with mild local tissue inflammation and no signs of systemic toxicity over the 20-month period. Overall, our refillable nISL implant is a promising ultra-long-acting delivery technology for HIV PrEP.

**One Sentence Summary:** An ultra-long-acting and subcutaneous refillable nanofluidic implant achieved preventive levels of islatravir in non-human primates for 20 months without refilling and conferred 100% protection against rectal and vaginal SHIV transmission.

## 1. Introduction

When taken as prescribed, oral single tablet combination of emtricitabine (FTC)/tenofovir disoproxil fumarate (TDF) or FTC/tenofovir alafenamide (TAF) are effective as pre-exposure prophylaxis (PrEP) against human immunodeficiency virus (HIV) transmission.(*1-6*) However, consistent adherence to daily dosing regimen is challenged by pill fatigue, forgetfulness, stigma or lifestyle discordance.(*7-10*) Moreover, less than 5% of at-risk individuals initiate PrEP(*11, 12*), further underscoring that effective uptake and implementation hinges on transformative changes in drug administration. To this end, long-acting approaches with infrequent dosing intervals, such as injectables, rings and implants, aim at improving therapeutic adherence, uptake, and persistence. In fact, long-acting ARVs that are discreet and offer long protection durations are likely viable alternatives for people at-risk for HIV infection.(*13-15*)

Long-acting ARV strategies typically focus on a single active pharmaceutical ingredient (API) instead of multi-drug formulations to maximize loading.(*16-25*) This allows for minimizing either injection volumes or implant size dimensions to reduce discomfort, pain or visibility. Ideally, the selected API for long-acting ARV must have high potency (at low doses) against multiple HIV strains as well as long-term stability, safety and tolerability. To date, cabotegravir, rilpivirine, TAF, dapivirine and islatravir are examples of prime API for long-acting approaches. Most recently, cabotegravir was approved as a long-acting injectable for HIV PrEP with a 2-month dosing frequency after two initial injections administered one month apart. Long-acting injectables often suffer from drawbacks of burst release, large injection volumes and site-specific adverse reactions, and risky year-long sub-therapeutic tails. Moreover, most injectables cannot be removed in the event of medical complications. On that note, dapivirine vaginal rings can be removed and afford discreet use(*26*). However, the need for monthly replacement poses adherence challenges especially in women under 25.(*27*) Numerous subdermal long-acting implants are under preclinical investigations for HIV PrEP(*25, 28-33*). Long-acting implants can circumvent high peaks and terminal dose dumping associated with injectables(*34*). Of these, TAF- and islatravir (ISL)-eluting and subdermal implants are under investigation(*35, 36*). Although TAF is a highly potent ARV, full protection against rectal SHIV challenges was not achieved with mono-PrEP in nonhuman primates (NHP)(*30, 37*).

ISL is a promising nucleoside reverse transcriptase translocation inhibitor that advanced to Phase 3 clinical trials as a single-agent ARV for HIV prevention. ISL has a long half-life (∼128.0 hrs)(*38*) and high barrier to drug resistance(*39*), and is ∼10-fold more potent than other ARV(*40*) with efficacy against multiple resistant HIV strains, rendering it ideal for single-agent LA PrEP(*40*). While Merck’s ISL-eluting implant has a projected release duration of 1-year, such polymeric eluting systems are associated with triphasic release kinetics and do not sustain constant drug delivery. Although once-weekly oral dosing of ISL is shown to be 100% effective in preventing rectal SHIV infection, PrEP efficacy of a ISL-releasing subdermal implant is yet to be determined.(*35, 41*)

In 2021, clinical trials with ISL for HIV treatment and PrEP were put on hold due to declines in CD4+ T cell or total lymphocyte counts, respectively.(*42*) These declines were determined to be dose-dependent, and specifically linked to high C_max_ levels associated with once-monthly oral dosing. In October 2022, Merck announced that while the study with once-monthly oral dosing was permanently ended, clinical trials using ISL at lower doses are resuming. Specifically, the Phase 2 clinical trial of once-weekly oral combination ARV therapy with lenacapavir and lower dose ISL will resume (NCT05052996). Similarly, a new Phase 3 clinical trial will be initiated for a once-daily oral combination of doravirine and a lower dose of ISL. No announcement was yet made regarding the clinical trial with ISL-eluting implants. However, it is plausible that the trial will resume given the favorable PK profile with low C_max_ levels associated with long-acting subcutaneous implants.

In light of these recent events, our ISL PrEP study is timely as we demonstrate long-term steady pharmacokinetics (PK) and complete protection for both rectal as well as vaginal infection, which was not shown previously. Here, we present an ultra-long-acting subcutaneous ISL delivery nanofluidic implant (nISL) for HIV PrEP. Our ultra-long-acting and transcutaneously refillable platform achieves constant ISL release for at least 20 months uninterrupted, which is unprecedented in HIV PrEP. We assessed the efficacy of sustained subcutaneous delivery of ISL via the nISL for protection from simian HIV_SF162P3_ (SHIV_SF162P3_) infection using repeated low-dose rectal or vaginal challenge in male and female NHP, respectively. We investigated the long-term PK and biodistribution of ISL. Lastly, we evaluated the safety and tolerability of the implant, as well as characterized the local foreign-body response to the implant.

## 2. Results

### 2.1. Refillable nanofluidic implant and transcutaneously injectable ISL formulation

To provide extended LA PrEP from a single drug-delivery device, we used a nanofluidic platform for sustained ISL release. Medical-grade titanium implants (20 mm × 13 mm × 4.5 mm, 570 μL reservoir volume) are equipped with two self-sealing silicone ports that provide access to the drug reservoir (Fig. 1A, top implant) for loading and refilling, as needed. A silicon nanofluidic membrane (6 mm × 6 mm × 0.4 mm) that harbors 278,600 nanochannels is mounted within the reservoir (Fig. 1B).(*43*) The nanofluidic membrane maintains sustained and steady drug elution via steric and electrostatic interactions between drug molecules and channel walls (Fig. 1A, bottom implant), wherein the nanochannel size and numbers as well as drug solubilization kinetics determine the release rate.(*44, 45*) Constant and sustained drug release is achieved passively across the nanofluidic membrane without the need for mechanical components or actuation.(*46, 47*) An external silicon carbide coating on the nanofluidic membrane ensures bioinertness and long term stability under physiological conditions. Figure 1C shows the integrity of a nanofluidic membrane after 4-month implantation in NHP.

**Fig. 1.**
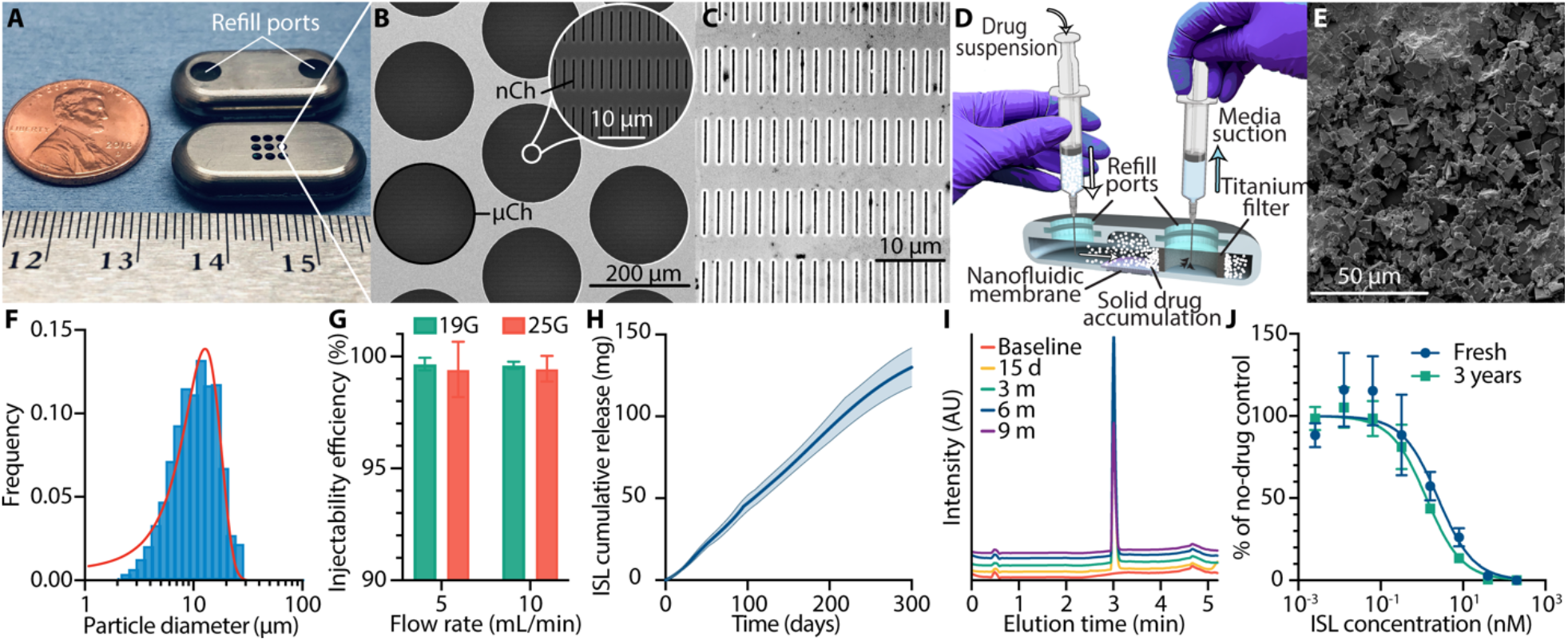
Refillable nanofluidic implant, ISL formulation for transcutaneous refilling, and in vitro ISL release. **(A)** Assembled refillable titanium implant showing the loading/refill ports side (top) and the drug-eluting membrane side (bottom). **(B)** SEM image of the silicon nanofluidic membrane showing circular cross section of microchannels (μCh) and inlet of slit-nanochannels (nCh) in the inset. **(C)** SEM of nanochannel membrane after 4 months of implantation in NHP. **(D)** Cross-section schematic of refillable nanofluidic implant demonstrating solid particle drug refilling. **(E)** SEM image of ISL particles. **(F)** ISL particle size distribution in PBS suspension with 0.05% Tween 20. **(G)** Injectability efficiency of ISL particles using 19G and 25G needles. **(H)** In vitro cumulative ISL release (mean ± SD) from nISL loaded with ISL suspension in PBS + 0.05% Tween 20 (n=5). **(I)** HPLC ISL chromatogram @261 nm from sink solution samples collected at baseline, day 15, and months 3, 6, 9. **(J)** TZM-bl infectivity assay showing similar bioactivity of ISL after solubilization in PBS and 3 years of incubation at 37 °C compared to fresh stock (mean ± SD).

The implant allows solid drug particle loading and refilling for maximum loading efficiency and minimal implant volume. For this, drugs are formulated in a microparticle suspension. Solid drug loading and refilling are accomplished using simple injection and venting syringes owing to the inclusion of a sintered titanium filter (4 mm inner diameter, 6 mm outer diameter, 2.2 mm height) under a silicone port (Fig. 1D). The titanium filter allows the suspension media to pass through, while solid particles are retained within the reservoir, resulting in solid drug loading.(*48, 49*) This innovative refillability feature of the implant affords extension of treatment duration indefinitely.

For solid drug loading and refilling of the implant, syringeability and injectability are key parameters. Thus, an ISL microparticle suspension was formulated using 0.05% Tween 20 in PBS to prevent needle clogging during suspension transfer from the vial and facilitate implant refilling.(*48*) Tween 20 is a commonly used excipient in FDA approved pharmaceutics.(*50*) Representative SEM image (Fig. 1E) shows ISL particles within the suspension. In the suspension, 95% of the ISL particles were within 1-19 μm (Fig. 1F). Using a medical grade titanium filter with 10 μm nominal porosity, we measured particle retention efficiency of 99.1%. We achieved ISL loading efficiency 89 ± 4.8% defined as the percentage of drug retained in the implant vs drug injected upon loading. Further, we measured ISL loading density of 0.85 mg/μL. Injectability efficiency >99%, defined as the percentage of drug injected vs drug leftover in the syringe, was obtained at injection rates of 5 or 10 mL/min and with 19G and 25G needles (Fig. 1G), indicating no sedimentation occurred. In vitro release studies at 37 °C using implants loaded with 150 mg ISL formulated in a suspension with 0.05% Tween 20 (nISL) (n=5) showed zero-order release in PBS (Fig. 1H). Linear regression of the cumulative release revealed an average release of 462.9 ± 2.9 μg/d. HPLC chromatogram of ISL released from the implants for up to 9 months (Fig. 1I) had no secondary peaks, demonstrating drug stability. TZM-bl cell infectivity assay using HIV-1NL4-3 demonstrated that the calculated half maximal effective concentration (EC_50_) of ISL incubated for 3 years in PBS solution at 37 °C and fresh ISL were 1.29 nM (95% CI, 0.99 to 1.67 nM) and 2.47 nM (95% CI, 1.39 to 4.41 nM), respectively (Fig. 1J). This supports ISL as a suitable ARV for long-term release from reservoir-based implantable devices.

### 2.2. nISL pharmacokinetic profile in NHP

For in vivo evaluation of the long-term PK profile, male rhesus macaques (n=4) were subcutaneously implanted, contralaterally in the dorsum, with two nanofluidic implants presenting 360 nm nanochannels, both initially filled with PBS (Fig. 2A). One of the four animals was added to the study 8 months later due to animal availability. Implants were oriented with membrane releasing toward the subcutaneous tissue, while the silicone port side was facing the epidermis, allowing access for transcutaneous loading and refilling. After implantation, one of the implants in each animal was immediately transcutaneously loaded with ISL suspension using the procedure described above and in our previous publication.(*51*) The loading port was identified visually and via palpation. Transcutaneous loading was executed within 60 seconds for each implant. Mechanistically, upon loading, interstitial fluids permeate across the nanochannel membrane into the implant reservoir and dissolve a portion of solid ISL particles. Solubilized drug molecules are then released from the implant by diffusion across the nanochannels, where their size and number control the release rate. This establishes a stable mechanism of drug solubilization and nanoconfined release that terminates only when the implant is completely depleted of drug.

**Fig. 2.**
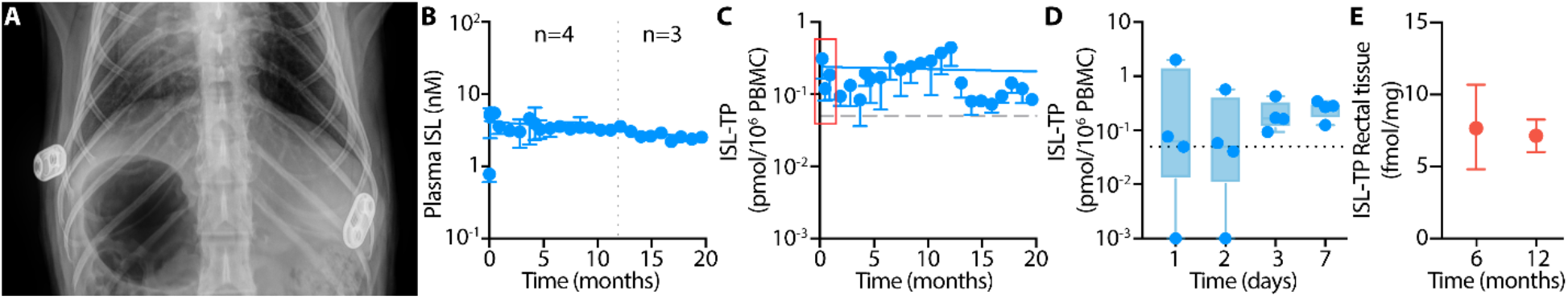
Pharmacokinetics and tissue distribution of ISL from NHP implanted with subcutaneous nISL. **(A)** X-ray image of NHP implanted with two contralateral nanofluidic devices. **(B)** 20 months of steady plasma ISL concentrations in NHP subcutaneously implanted with nISL. One animal was added to the study 8 months after the first cohort of 3 macaques (n=4 until month 12, n=3 thereafter). This animal is currently on month 12 of the pK study. **(C)** 20 months of PBMC intracellular ISL-TP concentrations in NHP subcutaneously implanted with nISL. Horizontal dashed gray line represents preventive threshold target of 0.05 pmol/10^6^ cells. Early time points in the red box are shown in panel D. **(D)** PBMC ISL-TP concentrations for each animal for the first 7 days of the study. **(E)** ISL-TP concentrations in rectal tissue biopsy at defined time points. All data are presented as median and IQR.

Plasma ISL concentrations were consistently maintained throughout the 20-month study (Fig. 2B) at 3.12 nM (Median, IQR, 2.61 to 4.05 nM; mean, 3.45 nM) indicating uninterrupted sustained release from nISL. These are similar to average plasma levels (2.8 nM) observed in the study of once-weekly oral ISL dosing (3.9 mg/kg) in NHP.(*52*) We used PBMC ISL-TP concentration of 0.05 pmol/10^6^ cells as the prevention target as previously reported.(*35, 52*) After a single transcutaneous loading of the implant with ISL, target preventive ISL-TP concentrations in PBMCs were surpassed and maintained at a median of 0.16 pmol/10^6^ cells (IQR, 0.086 to 0.30 pmol/10^6^ cells) for 20 months (Fig. 2C). The preventive threshold was surpassed by day 1 for 2 of the 4 animals (Fig. 2D). The other two animals achieved preventative levels by day 3. Further, we measured ISL-TP concentrations in rectal tissue biopsies, which is a relevant site for HIV-1 transmission or a key viral reservoir. Drug accumulation in rectal tissue was maintained at a median of 7.65 fmol/mg (IQR, 4.82 to 10.70 fmol/mg) and 7.14 fmol/mg (IQR, 6.00 to 8.25 fmol/mg) at 6 and 12 months, respectively (Fig. 2E). This suggests that ISL tissue biodistribution was achieved and maintained.

### 2.3. Preventive efficacy of nISL against rectal and vaginal SHIV_SF162P3_ infection

We next assessed whether sustained ISL delivery as a subcutaneously delivered monotherapy could protect the macaques against rectal and vaginal SHIV_SF162P3_ infection.

For the rectal challenge study, animals were subcutaneously implanted contralaterally in the dorsum with two PBS-filled nanofluidic implants presenting 280 nm nanochannels. One of the two implants was transcutaneously loaded with ISL immediately after implantation as previously described. Prior to SHIV challenge, the animals were subjected to a one-month “conditioning phase” (Fig. 3A) during which PBMC ISL-TP concentrations were monitored to ensure that the target preventive threshold of 0.05 pmol/10^6^ cells was achieved (Fig. 2C) prior to initiating the viral challenges. All animals with nISL (PrEP cohort) surpassed the preventative threshold by day 1 post-implantation. Animals in both nISL PrEP (n=6, male) and controls (n=12, 9 male and 3 female) cohorts were rectally challenged weekly with low-dose (4.27 TCID) SHIV_SF162P3_ for up to 10 inoculations (Figure 3A). Of the control cohort, six males served as ‘real-time’ controls, while six animals were historical controls (3 male and 3 female) from earlier studies performed under identical conditions with the same virus stock, inoculum size and inoculation intervals and period.(*30*) All animals were monitored for PK throughout the 6-month study inclusive of post-device explantation on week 16 for a drug washout phase.

**Fig. 3.**
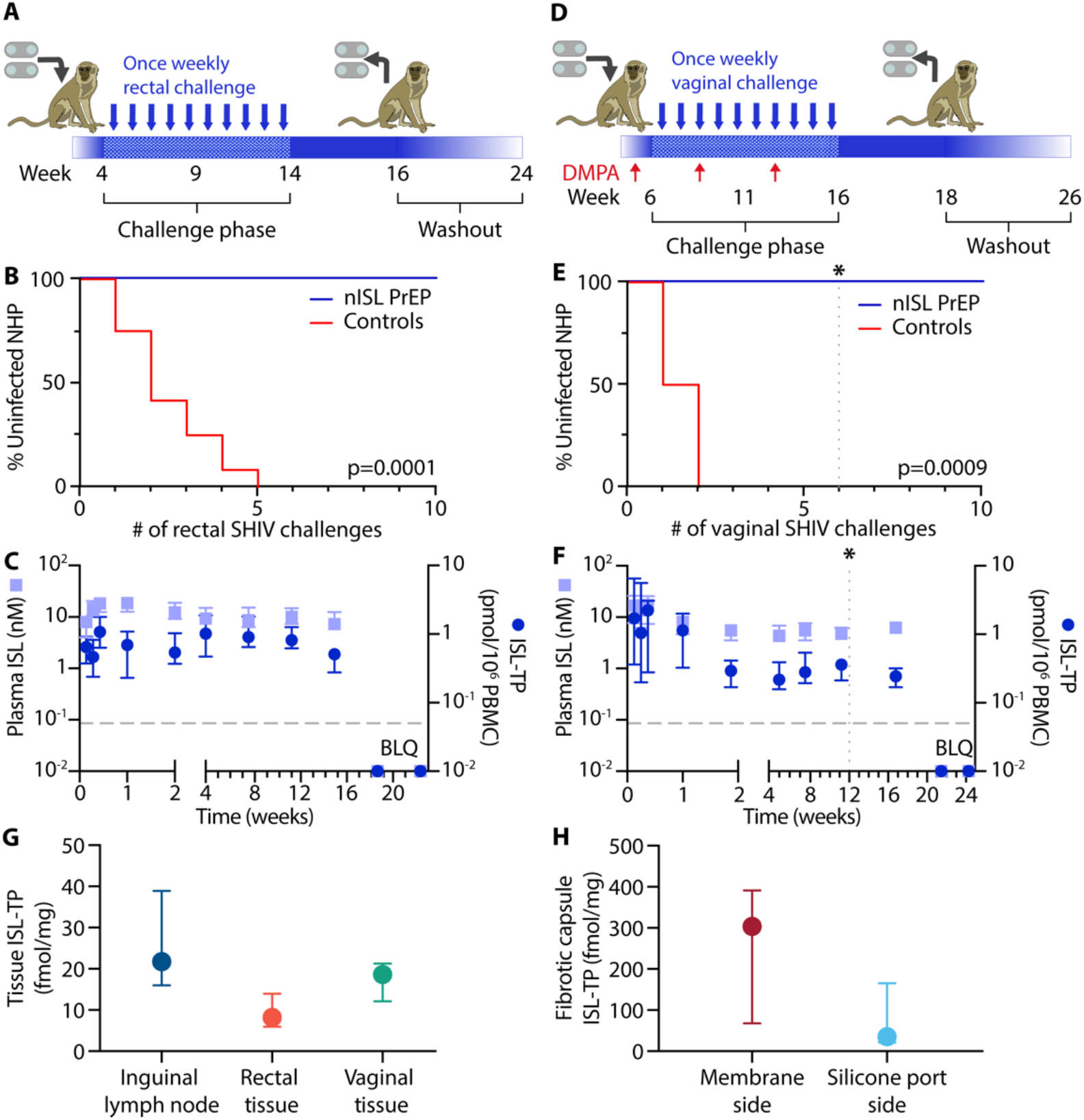
PrEP efficacy of nISL against rectal and vaginal SHIV. **(A)** Schematic of ISL PrEP efficacy study design for rectal SHIV challenge model. Conditioning phase to ensure PBMC ISL-TP concentrations above 0.05 pmol/10^6^ cells (0-4 weeks). Rectal challenge phase with up to 10 weekly low-dose (4.27 TCID) SHIV_SF162P3_ exposures. ISL PK continuation phase (week 14-16) followed by nISL explantation from all animals at week 16. ISL washout was observed for 8 weeks in which seronegative animals were monitored in the absence of drug. **(B)** Kaplan-Meier curve representing the percentage of uninfected animals as a function of weekly rectal SHIV exposure. PrEP (n=6, male) vs control (n=12, 9 male and 3 female) group. **(C)** ISL plasma and PBMC ISL-TP concentrations in PrEP group throughout rectal efficacy study. **(D)** Schematic of ISL PrEP efficacy study design for vaginal SHIV challenge model. Conditioning phase to reach PBMC ISL-TP concentrations above 0.05 pmol/10^6^ cells and depot medroxyprogesterone acetate (DMPA, DepoProvera) injection (30 mg) 2 weeks before SHIV challenges. Thereafter, DMPA injections were administered monthly in seronegative NHP during challenges. Vaginal challenge phase with up to 10 weekly low-dose (50 TCID) SHIV_SF162P3_ exposures. ISL PK continuation phase followed by nISL explantation from all animals at week 16. ISL washout was observed for 8 weeks in which seronegative animals were monitored in the absence of drug. **(E)** Kaplan-Meier curve representing the percentage of uninfected animals as a function of weekly vaginal SHIV exposure. PrEP (female, n=6 up to challenge week 6(*); due to implant loss, one animal was censored thereafter) vs control (n=6, female) group. **(F)** Plasma ISL and PBMC ISL-TP concentrations in PrEP group throughout the vaginal efficacy study (n=6 up to study week 10, corresponding to challenge week 6(*); n=5 thereafter due to censored animal). **(G)** Inguinal lymph node (ILN), rectal, and vaginal tissue ISL-TP concentrations after explantation for the vaginal PrEP study. **(H)** Fibrotic capsule ISL-TP concentrations near membrane and ports for the vaginal PrEP study. Survival curve statistical analysis by Mantel-Cox test. (*) One animal censored. Data presented as median and IQR. (* = p<0.05)

To monitor for SHIV_SF162P3_ infection, we evaluated cell-free plasma viral RNA levels weekly. Rectal challenges were stopped upon initial detection of plasma viral RNA, which was confirmed after a consecutive positive assay. The six real-time controls were infected at challenges 1 (n=1), 2 (n=3), 3 (n=1) and 5 (n=1), with a median time to infection of 2 challenges. All six macaques from the nISL PrEP group (6 out of 6, 100%) were uninfected after 10 weekly rectal SHIV_SF162P3_ challenges (Fig. 3B). The nISL PrEP cohort had a 12.90-fold lower risk of infection (95% CI of HR, 3.54-46.96; p=0.0001) compared to control. Kaplan-Meier analysis demonstrated statistical significance (p=0.0005) between nISL and control groups. After device explantation at month 4, there was no spike in viremia, indicative of PrEP efficacy of nISL monotherapy in the six uninfected animals.

Animals in the nISL PrEP cohort maintained a median plasma ISL concentration of 11.08 nM (IQR, 7.96 to 15.36 nM; mean ± SD, 12.33 ± 5.38 nM) for 4 months (Fig. 3C). In addition, median PBMC ISL-TP concentration of 0.66 pmol/10^6^ cells (IQR, 0.49 to 1.01 pmol/10^6^ cells; mean ± SD, 0.81 ± 0.52 pmol/10^6^ cells) were maintained for 4 months.

These levels were nearly 4-folds higher than observed during the long-term PK analysis which is consistent with the difference in animal weight for the two animal cohorts: median 11.01 kg (IQR 9.68 to 12.79 kg) for long-term PK vs 3.25 kg (IQR 3.17 to 3.37 kg) for the rectal challenges study.

These levels were identical to the PBMC ISL-TP levels (mean, 0.81 pmol/10^6^ cells) observed at the time of intrarectal challenge in NHP receiving once-weekly oral ISL dosing (3.9 mg/kg), which conferred full protection.(*53*)

The vaginal challenge study was performed in a like manner to the rectal experiment (Fig. 3D) with nISL PrEP (n = 6, female) and control (n = 6, female) cohorts. During the one-month “conditioning phase”, all animals surpassed the preventative target by day 1 post-implantation (Supplementary Materials Fig. S1). Thereafter, all animals received an injection of depot medroxyprogesterone acetate (DMPA, Depo-Provera) to thin the vaginal epithelium 2 weeks prior to the start of SHIV challenge. DMPA dosing was performed once every four weeks in seronegative animals during challenges to ensure progesterone levels remained suppressed.(*54*) Animals in both PrEP (n=6) and control (n=6) cohorts were vaginally challenged weekly with low-dose (50 TCID) SHIV_SF162P3_ for up to 10 inoculations and continually monitored for drug PK throughout the 6-month study (Figure 3D).

Vaginal challenges were stopped upon detection of plasma viral RNA, which was confirmed after a consecutive positive assay. The six real-time controls were infected at challenges 1 (n=3) and 2 (n=3) (Fig. 3E). Animals in the nISL PrEP group with detectable PBMC ISL-TP concentration (n = 5) were uninfected after 10 weekly vaginal SHIV_SF162P3_ challenges (Fig. 3E). One animal in the nISL PrEP group was infected at challenge 6 on week 12. Upon further investigation, we noted that this animal had lost the implant and had undetectable plasma ISL and PBMC ISL-TP concentrations on week 12. On week 8, this animal had preventive PBMC ISL-TP concentrations. Thus, we surmise that the loss of nISL implant occurred thereafter. While rare, the loss of an implant can occur with the animal removing and discarding the device. In these cases, skin lesions from self-removal heal rapidly in healthy animals and can be concealed by fur. The animal was censored due to lack of nISL during the challenge phase. In this vaginal challenge study, the nISL PrEP group had a 23.17-fold lower risk of infection (95% CI of HR, 3.62-148.44; P=0.0009) compared to control. Animals remained seronegative after nISL explantation at month 4, indicative of PrEP efficacy of ISL monotherapy against vaginal transmission.

Macaques in the vaginal challenge cohort maintained a median plasma ISL concentration of 6.16 nM (IQR, 4.15 to 7.77 nM; mean ± SD, 7.79 ± 5.84 nM) for 4 months (Fig. 3F). Notably, these levels were substantially lower than measured in male macaques even when accounting for differences in animal weight (Median 4.27 kg, IQR 3.87 to 4.58 kg), which indicates sex-related differences in ISL PK. During this time, PBMC ISL-TP concentrations were maintained at a median of 0.33 pmol/10^6^ cells (IQR, 0.19 to 0.53 pmol/10^6^ cells; mean ± SD, 0.62 ± 0.82 pmol/10^6^ cells). Importantly, this data establishes a level of PBMC ISL-TP concentration that is protective against vaginal SHIV transmission, which was previously unknown.

At the time of device explantation, ISL-TP concentrations in inguinal lymph node (ILN), rectal and vaginal tissue biopsies were 21.9 fmol/mg (IQR, 16.1 to 39.0 fmol/mg), 8.24 fmol/mg (IQR, 6.0 to 14.0 fmol/mg) and 18.6 fmol/mg (IQR, 12.2 to 21.3 fmol/mg), respectively (Fig. 3G). ILN showed statistically significant higher levels of ISL-TP than both rectal (p = 0.01) and vaginal (p = 0.047) tissue. No statistical significance was observed between rectal and vaginal tissue (p = 0.11). ISL-TP concentration in the fibrotic capsule (FC) in contact with the membrane was higher (304.0 fmol/mg; IQR, 68.3 to 391.5 fmol/mg) than that of the silicone port side (35.4 fmol/mg (IQR, 21.3 to 165.3 fmol/mg) (Fig. 3H). However, there was no statistical significance between the two groups (p = 0.318).

Drug washout was monitored for 2 months after nISL explantation from all seronegative animals (6 male and 5 female macaques). During the washout period, both plasma ISL and PBMC ISL-TP concentrations were below the limit of quantitation (BLOQ) 1-month post-device retrieval for all animals (Fig. 3C, F). Further, we assessed potential presence of SHIV in various tissues biopsied from the nISL PrEP cohort by measuring cell-associated SHIV_SF162P3_ provirus DNA. All tissues were negative for SHIV in the PrEP cohorts in both rectal and vaginal challenge studies (data not shown).

To evaluate nISL release rate in vivo, residual contents from the implant were extracted and analyzed for ISL (Table 1). The nISL implants in the rectal and vaginal PrEP efficacy study each had a mean release rate of 0.52 ± 0.05 mg/day and 0.50 ± 0.25 mg/day, which was sufficient to sustain PBMC ISL-TP concentrations above 0.05 pmol/10^6^ cells throughout the duration of the study (Table 1).

**Table 1.** Residual drug analysis from nISL implants at explantation via high performance liquid chromatography (HPLC). Data presented as mean ± SD.

| NHP study | ISL loaded (mg) | Residual ISL (mg) | ISL release rate (mg/day) |
| --- | --- | --- | --- |
| Rectal (Male) | 187.09 ± 9.35 | 125.60 ± 6.39 | 0.52 ± 0.05 |
| Vaginal (Female) | 181.46 ± 4.94 | 117.40 ± 28.33 | 0.50 ± 0.25 |

### 2.4. nISL safety and tolerability in NHP

For the animals in the long-term PK study, nISL and PBS implants were generally well tolerated without systemic effects, deaths, or serious adverse events (SAE). The long-term PK analysis in NHP is currently ongoing. The study will be continued for an additional 16 months to assess long-term deployment (for a total of 36 months) and efficacy of transcutaneous refilling at 24 months post-implantation. Therefore, histopathology assessment of this cohort is not possible. To date, close monitoring revealed no signs of inflammation, erythema or discomfort to the animals. An exception was that one animal showed sudden onset of local tissue inflammation around the implant at 20 months post-implantation. There were no prior signs of swelling or inflammation. Upon surgical retrieval, we noted that the implant had flipped, with the membrane side oriented toward the epidermis instead. While the circumstances of the implant flipping are unclear, it could be attributable to animal manipulation of the implantation site. Therefore, we postulate that the sudden onset of local tissue inflammation could be due to release of ISL toward the skin as well as potential local manipulation of the implant by the animal. For this animal, histopathology analysis of the tissues surrounding the implant are provided in Supplementary Materials (Fig. S2).

Similar to the long-term PK study, for the rectal and vaginal challenge PrEP efficacy studies, nISL and PBS implants were well tolerated without systemic effects, deaths, SAE or discontinuation throughout the 16 weeks of implantation (Supplementary Materials Table S1). None of the animals had SAE, although 83% of the rectal nISL and 60% of vaginal nISL cohorts experienced transient moderate swelling or induration at the implant site.

To assess tissue reaction at the implantation site, we evaluated the nISL and contralateral PBS implants from the PrEP animals (Fig. 4A-D). The change in implantation site thickness (ΔT) with respect to depth (z-axis) was normalized to the baseline measurement taken on day of implantation (Fig. 4E). Rectal PBS control (R-PBS), rectal nISL (R-nISL), vaginal PBS control (V-PBS) and vaginal nISL (V-nISL) groups maintained a mean depth ΔT of 0.27 ± 0.45, 1.32 ± 0.82, 0.43 ± 0.51 and 1.45 ± 0.67 mm, respectively, over 4 months (Fig. 4F-I). Statistical significance was observed between the overall means of R-PBS and R-nISL (p=0.0023) and V-PBS and V-nISL (p=0.0042) (Fig. 4J). Similarly, the change in implant site thickness (ΔT) with respect to length and width (x-y plane) was normalized to the baseline measurement (Fig. 4E). R-PBS, R-nISL, V-PBS and V-nISL groups maintained a mean x-y plane ΔT of -0.22 ± 0.73, 0.95 ± 1.56, 1.00 ± 0.57 and 2.13 ± 1.88 mm, respectively, over 4 months (Fig. 4K-N). Statistical significance was observed between the overall means of R-PBS and R-nISL groups (p=0.045) (Fig. 4O).

**Fig. 4.**
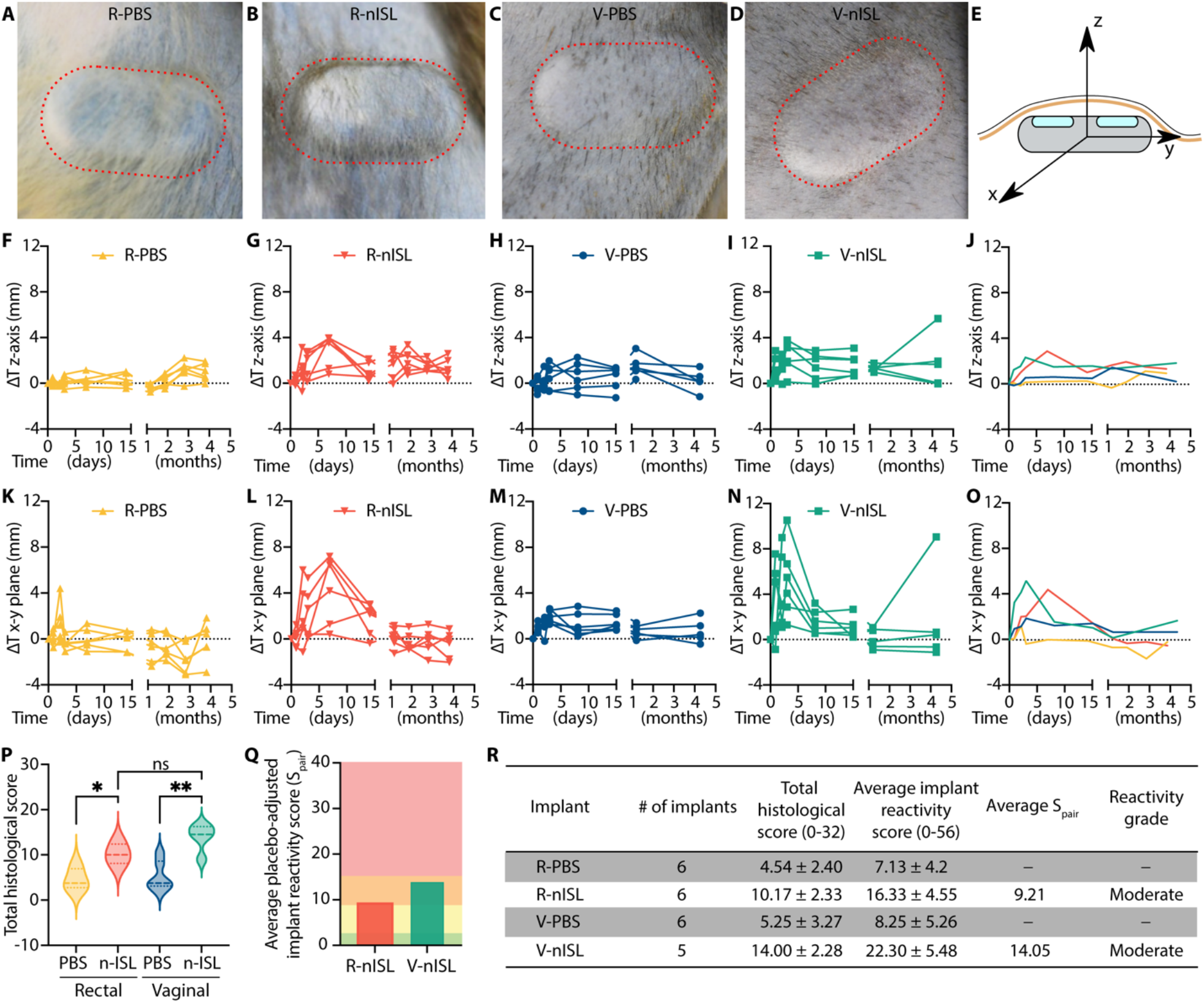
Safety and tolerability of nISL and PBS implants. Representative image of **(A)** rectal control PBS (R-PBS), **(B)** rectal nISL (R-nISL), **(C)** vaginal control PBS (V-PBS), **(D)** vaginal nISL(V-nISL) in NHP at 4 months of subcutaneous implantation. **(E)** Subcutaneous implant schematic demonstrating x, y and z axes. Change in implantation site thickness (ΔT) with respect to depth (z-axis) throughout 4 months in **(F)** R-PBS, **(G)** R-nISL, **(H)** V-PBS, **(I)** V-nISL, **(J)** comparison of 4 groups. Change in implant thickness (ΔT) with respect to length and width (x-y plane) throughout 4 months in **(K)** R-PBS, **(L)** R-nISL, **(M)** V-PBS, **(N)** V-nISL, **(O)** comparison of 4 groups. **(P)** Comparison of total histological scores between R-PBS, R-nISL, V-PBS and V-nISL groups. **(Q)** Comparison of average S_pair_ reactivity grade between R-nISL and V-nISL. S_pair_ values: 0.0-2.9, 3.0-8.9, 9.0-15.0, and >15.1 colored as green (no reaction), yellow (slight reaction), orange (moderate reaction) and red (severe reaction). **(R)** Table with histopathological scoring in all 4 groups.

Two blinded board-certified pathologists scored the tissue directly in contact with the implant to assess foreign-body reaction and tolerability. After 4 months of subcutaneous implantation, the total histological score (scale 0 to 32) was 4.54 ± 2.40, 10.17 ± 2.34, 5.25 ± 3.27, and 14.00 ± 2.98 in the R-PBS, R-nISL, V-PBS and V-nISL groups, respectively (Fig. 4P). Likewise, the average implant reactivity scores were 7.13 ± 4.20, 16.33 ± 4.55, 8.25 ± 5.26, and 22.3 ± 5.48, respectively. There was statistical significance between the R-PBS group with respect to R-nISL (p=0.0012) and V-PBS with respect to V-nISL (p=0.0066). The average placebo-adjusted implant reactivity scores (S_pair_) for R-nISL and V-nISL was 9.21 and 14.05, respectively (Fig. 4Q). In summary, tissue response to our nISL implants (average release rate 1.00 ± 0.40 mg/day (R-nISL) and 0.99 ± 0.24 mg/day (V-nISL)) was qualified as moderate reaction for both R-nISL and V-nISL (Figures 4P-R).

For safety and toxicity evaluation, we assessed kidney and liver enzymes and total lymphocyte count in the animals with nISL implants from the PK study as well as the rectal and vaginal challenge cohorts (Fig. 5A-I). Creatinine levels were within normal range for all studies, suggesting that there were no detectable kidney toxicities in the nISL cohorts (Fig. 5A, F). In addition, liver enzymes, aspartate aminotransferase (AST) (Fig. 5B, G) and alanine aminotransferase (ALT) (Fig. 5C, H), were within normal levels with respect to reference rhesus macaque levels. Moreover, no appreciable changes in AST or ALT were observed with respect to their baseline levels, which were measured prior to implantation.

**Fig 5.**
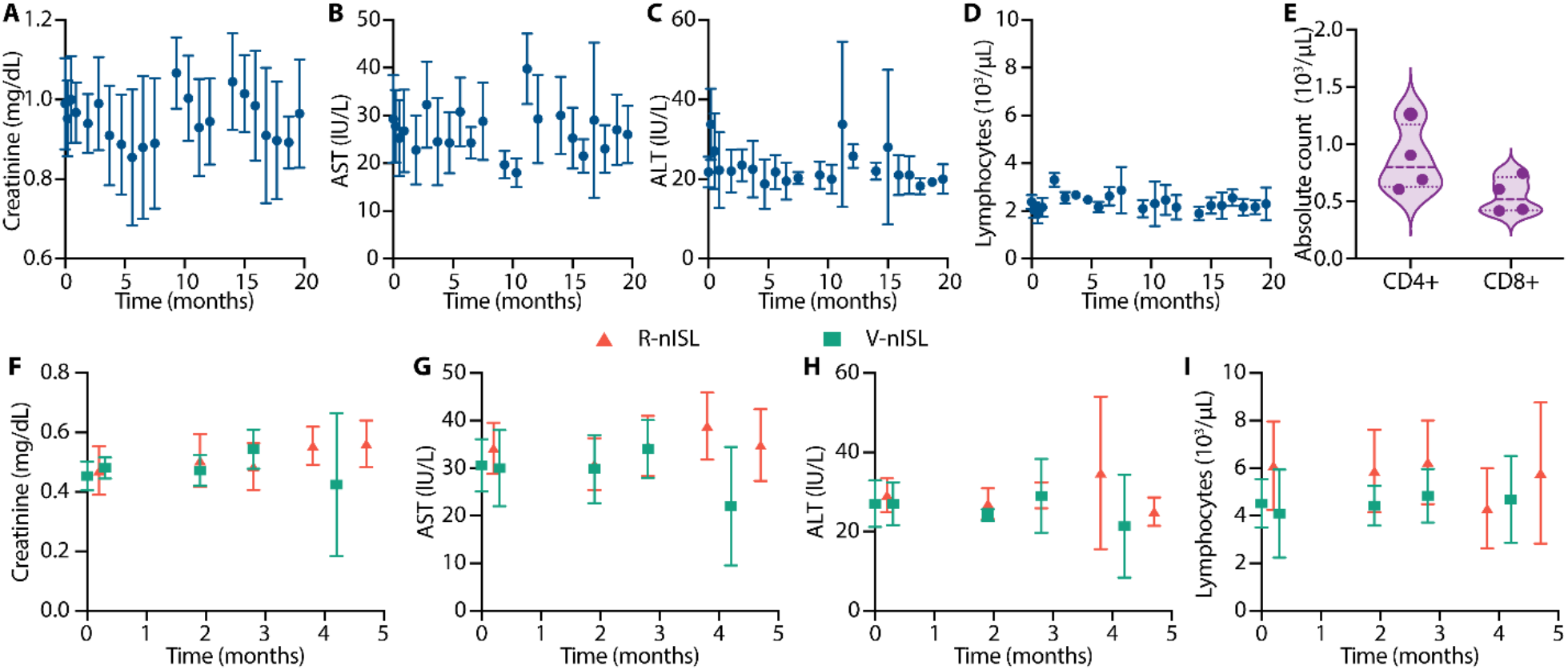
Liver and kidney enzymes and total lymphocyte levels. Longitudinal analysis of (A) creatinine activity, liver enzymes **(B)** aspartate aminotransferase (AST), and **(C)** alanine aminotransferase (ALT), **(D)** total lymphocyte count for the nISL PK study. **(E)** CD4+ and CD8+ T cells count from the PK study at month 18. Corresponding **(F)** creatinine, **(G)** AST, **(H)** ALT, **(I)** total lymphocyte count for animals with nISL implants in the rectal (red) and vaginal (green) SHIV challenge studies, respectively. Data presented as mean ± SD.

Total lymphocyte count remained stable for all cohorts (Fig. 5D, I). Notably, no changes with respect to baseline were detected throughout the 20-month PK analysis (Fig. 5D), with a median total lymphocyte count of 2120/μL (IQR, 1863 to 2513). CD4+ and CD8+ T cell counts were also assessed during the PK study on month 18. Median CD4+ and CD8+ T cell counts were 797/μL (IQR, 627 to 1172) and 518/μL (IQR, 421 to 711), respectively (Fig. 5E). While baseline was not quantified for these animals, these levels were within normal reference values for rhesus macaques.

Collectively, these are crucial findings given the temporary suspension of ISL clinical studies due to the dose-dependent decrease in total lymphocytes and CD4+ T cell count in trial participants.(*42*) In addition, metabolic panel and complete blood count results were also within normal levels in all three studies (Figures S3-S8 Supplementary Materials).

### 2.5. Characterization of foreign-body response to nISL in NHP

To assess foreign-body response to nISL, we histologically examined the tissue surrounding the implants after 4 months of implantation. Given that fibrotic capsules could potentially impede drug release from implants, we performed a deeper tissue characterization by investigating thickness, collagen density, and blood vessel number. These parameters could affect molecular transport locally. Further, we quantified effective diffusivity in fibrotic capsules surrounding the implants via fluorescence recovery after photobleaching (FRAP) to assess molecular transport.

The fibrotic capsules were examined via Masson’s Trichrome staining (Fig. 6A-D); median measurements were 104 μm (IQR, 80 to 120 μm), 479 μm (IQR, 276 to 578 μm), 159 μm (IQR, 114 to 214 μm) and 704 μm (IQR, 369 to 956 μm) for R-PBS, R-nISL, V-PBS and V-nISL, respectively. Statistical significance was noted for fibrotic capsule thickness between R-PBS and R-nISL (p<0.0001) and V-PBS and V-nISL (p<0.0001) (Fig. 6E).

**Fig. 6.**
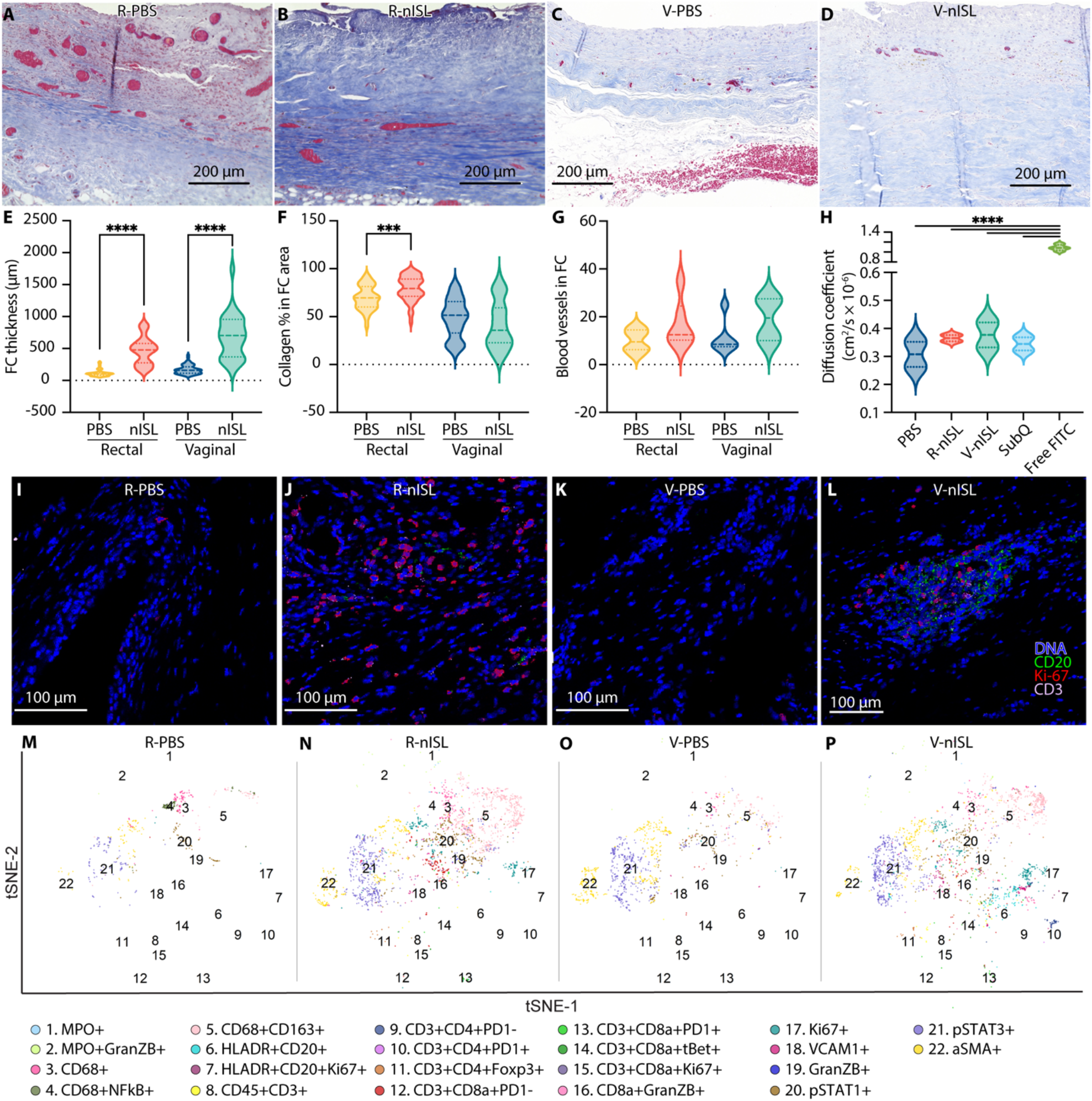
Characterization of fibrotic capsule surrounding nPBS and nISL. Masson’s Trichrome staining of 4-month fibrotic capsule (FC) in **(A)** rectal control PBS (R-PBS), **(B)** rectal nISL (R-nISL), **(C)** vaginal control PBS (V-PBS) and **(D)** vaginal nISL (V-nISL). Comparison of **(E)** FC thickness, **(F)** collagen percentage in FC area, **(G)** blood vessel count in FC and **(H)** FRAP diffusivity coefficient. Imaging Mass Cytometry (IMC) representative images of **(I)** R-PBS, **(J)** R-nISL, **(K)** V-PBS and **(L)** V-nISL fibrotic capsules. IMC t-SNE plots of representative images of **(M)** R-PBS, **(N)** R-nISL, **(O)** V-PBS and **(P)** V-nISL fibrotic capsules. Data presented as median ± IQR.

Collagen density percentage in the fibrotic capsule was evaluated in the Masson’s Trichrome stained samples; median measurements were 69.5 % (IQR, 60.0 to 81.5 %), 79.4 % (IQR, 71.2 to 89.2 %), 51.5% (IQR, 32.9 to 65.7 %) and 35.8% (IQR, 22.7 to 59.2 %) for R-PBS, R-nISL, V-PBS and V-nISL, respectively. Statistical significance was noted for collagen density percentage in the fibrotic capsule between R-PBS and R-nISL (p=0.0009) (Figure 6F). Moreover, median number of blood vessels in R-PBS, R-nISL, V-PBS and V-nISL was 9.5 (IQR, 6.3 to 14.5; mean ± SD, 10.0 ± 4.7), 12.5 (IQR, 10.3 to 24.5; mean ± SD, 16.7 ± 10.0), 8.5 (IQR, 7.5 to 14.5; mean ± SD, 11.2 ± 7.0) and 19.5 (IQR, 10.0 to 27.5; mean ± SD, 19.0 ± 9.4), respectively. No statistical significance was observed between R-PBS and R-nISL (p=0.17) and between V-PBS and V-nISL (p=0.17).

FRAP was performed on fibrotic capsules retrieved from V-PBS, V-nISL and R-nISL groups to determine the diffusion coefficient. We used fluorescein isothiocyanate (FITC), a small hydrophilic molecule, as a surrogate for ISL. Mean values were 3.08 ± 0.63 ×10^−6^ cm^2^/s, 3.77 ± 0.63 ×10^−6^ cm^2^/s and 3.65 ± 0.16 ×10^−6^ cm^2^/s for V-PBS, V-nISL and R-nISL, respectively (Fig. 6H). These values were compared to diffusivities in subcutaneous tissue (3.45 ± 0.33 ×10^−6^ cm^2^/s) and free FITC (10.88 ± 0.59 ×10^−6^ cm^2^/s) reported in Fig. 6H.

Overall, despite the thicker fibrotic capsules in the nISL groups, we observed similar collagen density, number of blood vessels and FITC effective diffusion coefficient to the PBS implant controls. These results indicate that the fibrotic capsule surrounding the implant nor its thickness have a significant impact on the release and local transport of hydrophilic small molecule drugs such as ISL.

The nature and frequency of subpopulations of immune cells in close proximity to the implant and production of inflammatory cytokines was also investigated. The local immune response in the fibrotic capsule surrounding the devices was characterized via imaging mass cytometry (IMC) analysis of R-PBS, R-nISL, V-PBS and V-nISL groups (Figures 6I-P, Supplementary Materials Figure S9, S10) and production of proinflammatory cytokines (Supplementary Materials Figure S11). All cohorts showed increased in local macrophage infiltration (population 3), aligning with typical foreign body response to implantable devices. In the nISL cohorts, there was an influx of pro-inflammatory macrophages (population 5), which could be induced by chronic local exposure to drugs. We observed an increase in regulatory T cells (population 5), which are immunosuppressive, in nISL cohorts. Further, even though there was a slight increase in granzyme B-releasing CD8+ T cells (population 16), we noted lack of activation or proliferative state (populations 12-15). The increase of VCAM1+ in nISL groups is suggestive of leukocytes recruitment to vasculature, in line with the augmented levels of aSMA (blood vessel marker). In summary, there was a fibrotic response to the implant, however minimal influx of activated inflammatory cells in the nISL samples aligned with moderate histological and reactivity scoring (Fig 4P-R). Bioplex analysis of fluids extracted from the implant reservoir at explantation confirmed this finding showing no differences in cytokine levels between control and nISL (Supplementary Material Figure S11). Interleukin 11 (IL-11) here acting as an anti-inflammatory cytokine showed elevated levels albeit without statistical significance.

## 3. Discussion

This study provides unprecedented advancement in long-acting ARV delivery systems for HIV PrEP, where uninterrupted drug delivery is sustained above the target preventive threshold for at least 20 months in vivo. Importantly, the nISL subcutaneous implant has favorable safety and tolerability profile and fully protects against rectal and vaginal SHIV infection. Considering that women are more vulnerable to HIV infection and need to maintain stricter adherence than men to maintain preventive drug levels when taking oral PrEP, efficacy against vaginal transmission is fundamental.(*55-58*) Of relevance, despite equivalent daily release rates, female macaques had nearly 45% lower plasma ISL and PBMC ISL-TP levels than males, which cannot be solely justified by the difference in animal weight. This result highlights sex-related differences in body-mass composition (i.e., muscle and fat), ISL PK, metabolism, and clearance, which will require further elucidation, particularly for clinical translation. Nevertheless, sex-related differences in PK are consistent with previous finding with various ARV drugs.(*59, 60*)

In light of this, our study supports that drug penetration in rectal and vaginal tissues are important for attaining full protection. ISL has equivalent penetration in rectal and vaginal tissues, in contrast to lower vaginal penetration observed with tenofovir disoproxil fumarate.(*61*) In line with this, tissue concentrations are important as HIV transmission could be blocked before it enters the systemic circulation. Nonetheless, this is the first report of ISL-TP tissue concentrations in a PrEP efficacy NHP model, where effective preventive levels are yet to be established.

Notably, our implantable ultra-long-acting ARV platform maintained median PBMC ISL-TP concentrations above the preventive benchmark of 0.05 pmol/10^6^ cells for 20 months after a single transcutaneous ISL loading. For 94% of animals, the preventive threshold was surpassed as early as one day after implantation. This is important in the context of clinical deployment as protection can be achieved with no significant delay post-implantation and initiation of PrEP. On this note, we reported similar PBMC ISL-TP concentrations as that achieved with weekly oral dosing of 3.9 mg/kg in NHP, which conferred full rectal protection.(*53*) Further, drug concentrations were undetectable soon after device retrieval. This highlights another advantage of retrievable implants, in contrast to LA injectables, which have long pharmaceutical tails that increases the risk of selecting for drug-resistant viruses(*62*).

Importantly, we demonstrated that total lymphocyte and CD4+ T cell counts were unaffected by long-term subcutaneous ISL administration. These results are opportune, considering the dose-dependent decreases observed in ISL human participants, which resulted in clinical trial holds. A plausible explanation is that the nISL releases at lower concentrations, compared to the high C_max_ achieved with once-monthly oral pill dosing in clinical trials. Although ISL metabolism differs between species,(*63*) this compelling finding motivates further investigations. Furthermore, the implant was well-tolerated, and few mild AEs were observed, such as implant site induration. Although these results were in NHP, they were similar to those reported in the safety and tolerability clinical trial for the polymeric ISL-releasing implant.(*35*) Moreover, the toxicity assessment demonstrated no kidney or liver damage in all three of the studies.

Characterizing the foreign-body response to a drug-releasing implant is vital for understanding its deployment lifespan. The fibrotic capsule immediately starts forming after implantation, where its characteristics depend on the drug-releasing implant.(*64*) In the context of small molecules such as ISL, our study demonstrated that drug diffusion is independent of fibrotic capsule thickness. This is a key result for drug-delivery technologies in which foreign body response is expected to occur. As anticipated, higher ISL-TP concentrations were found in the fibrotic capsule in contact with the membrane as opposed to the ports. Further, fibrotic capsule was thicker in the ISL-loaded implants than PBS controls, and the inflammatory response, albeit moderate, was confined to the side in corresponding to the direction of drug release.

The fibrotic capsule areas in proximity to the membrane were further investigated with IMC to gain insights into immune responses to sustained drug release. PBS-filled implants had minimal immune infiltration, whereas the limited influx of inflammatory cells, namely, neutrophils, T cells and macrophages observed for nISL is consistent with normal tissue response to implanted drug delivery devices. However, histopathology assessment of the nISL implant that unexpectedly flipped and released ISL toward the epidermis instead, presented a contrasting tissue response. In this case, chronic inflammation, necrosis, and presence of proliferative immune cells were indicative of a severe foreign body response (see Supplementary Materials Section 1, Fig. S2). This finding is noteworthy as it indicates that the directionality of drug release is a key factor in determining implant tolerability. We posit that for subdermal implants, drugs are most effectively cleared locally when released toward the subcutaneous tissue as opposed to the epidermis. The local accumulation of higher drug concentration within the epidermis, concurrent with potential external insults such as friction and mechanical stresses, can result in severe foreign body responses.

Most subcutaneous long-acting drug delivery platforms under development elute drugs into the surroundings through polymeric membranes that encase the entire implant and do not afford release directionality. Relevant to this, in our study of sustained subcutaneous TAF-eluting nanofluidic implants, we observed significantly milder foreign body responses than polymeric devices with an order of magnitude lower TAF release rate.(*30, 65, 66*) Overall, these results support our hypothesis that release directionality plays a fundamental role in implant tolerability. In terms of envisioned clinical deployment of our platform, implant flipping could be avoided by slight changes in aspect ratio to yield a slightly wider and thinner implant body. In a different context, the flat and wide aspect ratio of our implants likely contributes to preventing migration. In fact, no migration was observed in any of the animals. This is of relevance, considering that migrations remain a concern for other implantable technologies presenting thin cylindrical morphology.

In a clinical setting, drug-loaded implants can achieve similar PBMC concentrations as those transcutaneously filled after implantation.(*67*) Moreover, transcutaneous refillability is an attractive feature for extending drug release duration beyond 2 years. Refillability avoids the need for device replacement, which entail procedures that usually lead to pain and potential complications.(*68*) In fact, implants that are removable and have a release duration of at least 2 years are preferred by young women(*69, 70*), whereas biodegradability was ranked least important.(*70*) In some women, biodegradability raised concerns of where the implant goes in the body as it dissolves.(*71, 72*) Moreover, healthcare providers prefer palpability for removal, if necessary, which could also assure end-users that the implant has not migrated.(*73, 74*) However, palpability could be a challenge for discretion; thus, optimizing implant design and placement are key factors to consider for improving acceptability.

In conclusion, the nanofluidic implant is an ultra-long-acting refillable drug delivery platform that can deliver ISL in a sustained, uninterrupted manner for at least 20 months, and confer full protection against rectal and vaginal SHIV challenges. In light of recent human pharmacokinetic data related to polymeric ISL implants(*75*), our results could be rapidly extrapolated from NHP to humans. Overall, this work provides an ultra-long-acting PrEP alternative that could be adopted by all people at risk of HIV infection for improved uptake, adherence and implementation.

## 4. Experimental Section

### Nanofluidic implant assembly

Medical-grade 6AI4V titanium refillable drug reservoirs manufactured at the Machine Shop at Houston Methodist Research Institute. Briefly, a nanofluidic membrane possessing 278,600 nanochannels (mean; 280 nm) was mounted on the inside of the sterile drug reservoir as described previously.(*51*) Detailed information regarding the membrane structure and fabrication was described previously.(*47, 76*) Implants were welded together using Arc welding. Implants were primed for drug release through the nanofluidic membrane by placing implants in 1 X Phosphate Buffered Saline (PBS) under vacuum. This preparation method resulted in loading all implants with PBS. Implants were maintained in sterile 1X PBS in a hermetically sealed container and shipped for gamma sterilization at 30 kGy (VPTRad). Container with gamma sterilized implant in PBS was opened until implantation. ISL was purchased from MedChemExpress.

### ISL formulation characterization

Approximately 20 mg of ISL (MedChemExpress) was allocated in a vial with 2 mL of PBS with 0.05% Tween 20. Each vial was sonicated with a sonication probe at 50 amps for 4 min. Afterward 10 μL of drug solution was placed on a flat silica substrate onto an SEM holder and sputtered with 7 nm of iridium. SEM images were taken using FEI Nova NanoSEM 230 ultra-high resolution, at the HMRI Microscope Core. Another 10 μL of drug solution was placed on a microscope slide, using wedge slide technique. Slides were imaged at 40 × magnification using EVOS microscope on a 4 mm × 4 mm area (total 108 images per sample). Images were analyzed and particle size distribution determined using ImageJ and MATLAB.

### In vitro release from nanofluidic implant

The in vitro release study was performed using similar refillable resin nanofluidic devices. Devices (n=5) were assembled and refilled with ISL PBS with 0.05 % Tween 20 as previously described.(*51*) and placed in sink solution of 20 mL 1X PBS with constant agitation at 37°C. For analysis, the entire sink solution was retrieved and replaced with fresh PBS every other day for 300 days. The maximum ISL concentration regarding ISL saturation in sink solution was <10%, therefore maintaining sink condition. UV-Vis analysis was performed on a Beckman Coulter DU® 730 system. Absorbance was measured at 261 nm.

### Animals and animal care

All animal procedures were conducted at the AAALAC-I accredited Michale E. Keeling Center for Comparative Medicine and Research, The University of Texas MD Anderson Cancer Center (UTMDACC), Bastrop, TX. All animal experiments were carried out according to the provisions of the Animal Welfare Act, PHS Animal Welfare Policy, and the principles of the NIH Guide for the Care and Use of Laboratory Animals. All procedures were approved by the Institutional Animal Care and Use Committee (IACUC) at UTMDACC, which has an Animal Welfare Assurance on file with the Office of Laboratory Animal Welfare. IACUC #00001749-RN00. Indian rhesus macaques (*Macaca mulatta*; n=28; 16 males and 12 females) of 2-6 years and 3-16 kg bred at this facility were used in the study. All procedures were performed under anesthesia with ketamine (10 mg/kg, intramuscular) and isoflurane.

All animals had access to clean, fresh water at all times and a standard laboratory diet. Prior to the initiation of virus inoculations, compatible macaques were pair-housed. Once inoculations were initiated, the macaques were separated into single housing (while permitting eye contact) to prevent the possibility of SHIV transmission between the macaques. Euthanasia of the macaques was accomplished in a humane manner (IV pentobarbital) by techniques recommended by the American Veterinary Medical Association Guidelines on Euthanasia. The senior medical veterinarian verified successful euthanasia by the lack of a heartbeat and respiration.

### Implantation procedure

An approximately 1-cm dorsal skin incision was made on the left and right lateral side of the thoracic spine. Blunt dissection was used to make a subcutaneous pocket ventrally about 5 cm deep. The implants were placed into the pocket with the membrane facing the body. A simple interrupted tacking suture of 4-0 polydioxanone (PDS) was placed in the subcutaneous tissue to help close the dead space and continued intradermally to close the skin. All animals received a single 50,000 U/kg perioperative penicillin G benzathine/penicillin G procaine (Combi-Pen) injection and subcutaneous once-daily meloxicam (0.2 mg/kg on day 1 and 0.1 mg/kg on days 2 and 3) for postsurgical pain. Immediately post-implantation one of the two implants was manually refilled with their respective drug vial (ISL) using two sterile 5 mL syringes. The manual refill procedure consisted in pulling the plunger of the suctioning syringe while holding in place, without pushing, the loading syringe. The procedure stopped when all the filtered suspension media was collected in the suction syringe. Further details regarding the implant drug loading procedure can be found elsewhere.(*67*)

### Blood collection and plasma and PBMC sample preparation

All animals had weekly blood draws to assess plasma ISL concentrations, intracellular PBMC ISL-TP concentrations, plasma viral RNA levels, and cell-associated SHIV DNA in PBMCs. Blood collection and sample preparation were performed as previously described.(*23*) Blood was collected in EDTA-coated vacutainer tubes before implantation; on days 1, 2, 3, 7, 10, and 14; and then once weekly until euthanasia. Plasma was separated from blood by centrifugation at 1200 × *g* for 10 min at 4 °C and stored at -80 °C until analysis. The remaining blood was used for PBMC separation by standard Ficoll-Hypaque centrifugation. Cell viability was > 95%. After cells were counted, they were pelleted by centrifugation at 400 × *g* for 10 min, resuspended in 500 μL of cold 70% methanol/30% water, and stored at -80 °C until further use.

### Pharmacokinetic analysis of ISL in plasma and ISL-TP in PBMC and tissues

The PK profiles of ISL in plasma and ISL-TP in PBMC were evaluated throughout the 20 months in the long-term ISL PK study and throughout 4 months in the efficacy studies. Non-compartmental methods were used to summarize observed concentrations. After device explantation, drug washout was assessed for an additional 2 months in the efficacy studies (n=11).

All bioanalyses were performed at the CAVP laboratory at University of Colorado Anschutz Medical Campus. A reversed-phase ultra-performance liquid chromatographic tandem mass spectrometry (UPLC-MS/MS) assay for the determination of ISL in NHP plasma was developed. The method utilizes a stable labeled internal standard for ISL and a liquid-liquid extraction. The assay has a quantifiable range of 0.025 ng/mL to 100 ng/mL when 0.2 mL of plasma is analyzed. Intracellular ISL-TP concentrations in PBMC lysate and tissue homogenate were quantified using solid phase extraction and reversed-phase ultra-performance liquid chromatographic tandem mass spectrometry (UPLC-MS/MS) assay similar to that previously described.(*77*) Typically, 10 fmol/sample was used as the lower limit of quantitation (LLOQ). If additional sensitivity was needed, standards and quality controls were added down to 5 fmol/sample. Tissues (50-to 75-mg aliquots) were snap-frozen and stored at -80 °C until analysis.

### PrEP nISL efficacy against rectal and vaginal SHIV challenge

To study the efficacy of the PrEP implant against SHIV transmission, animals were divided into two groups, PrEP nISL-treated [n=12; 6 male (M) and 6 female (F)] or control nPBS (n=12; 6 M and 6 F), in a non-blinded study. The PrEP regimen consisted of subcutaneously implanted nISL for sustained drug release over 112 days. The efficacy of nISL in preventing rectal SHIV transmission was evaluated using a repeat low-dose exposure model described previously in the male macaques.(*78-80*) Six historical controls were added to the rectal PrEP efficacy survival curve since the same virus stock, dose and inoculation was used. In addition, the efficacy of nISL in preventing vaginal SHIV transmission was evaluated using a repeat low-dose exposure model. Moreover, a 30 mg DMPA injection was given 2 weeks prior to the start of challenges and monthly thereafter. (*81-83*) Animals were considered protected if they remained seronegative for SHIV RNA throughout the study. Briefly, a month after PrEP-treated macaques achieved PBMC ISL-TP concentrations above 0.05 pmol/10^6^ cells, groups were rectally or vaginally exposed to SHIV_SF162P3_ once a week for up to 10 weeks until infection was confirmed by two consecutive positive plasma viral RNA levels. The SHIV_SF162P3_ dose was in range of HIV-1 RNA levels found in human semen during acute viremia.(*80*)

Challenge stocks of SHIV_162p3_ were generously supplied by Dr. Nancy Miller, Division of AIDS, NIAID, through Quality Biological (QBI), under Contract No. HHSN272201100023C to the Vaccine Research Program, Division of AIDS, NIAID. The stock SHIV_162p3_ R922 derived harvest 4 dated 9/16/2016 (p27 content 173.33 ng/ml, viral RNA load >10^9^ copies/ml, TCID50/ml in rhesus PBMC 1280) was diluted 1:300 and 1:25.6 and 1ml of virus was used for rectal and vaginal challenge, respectively, each time.

For the challenge, the animals were positioned in prone position and virus was inoculated approximately 4 cm into the rectum or vagina. Inoculated animals were maintained in the prone position with the perineum elevated for 20 minutes to ensure that virus did not leak out. Care was also taken to not abrade the mucosal surface of the rectum and vagina.

### Infection monitoring by SHIV RNA in plasma and SHIV DNA in tissues

Infection was monitored by the detection of SHIV RNA in plasma using previously described methods(*84, 85*) with modification. Viral RNA (vRNA) was isolated from blood plasma using the Qiagen QIAmp UltraSense Virus Kit (Qiagen #53704) in accordance with manufacturer’s instructions for 0.5 mL of plasma. vRNA levels were determined by quantitative real-time PCR (qRT-PCR) using Applied Biosystems™ TaqMan™ Fast Virus 1-Step Master Mix (Thermofisher #4444432) and a primer-probe combination recognizing a conserved region of gag (GAG5f: 5’-ACTTTCGGTCTTAGCTCCATTAGTG-3’; GAG3r: 5’-TTTTGCTTCCTCAGTGTGTTTCA-3’; and GAG1tq: FAM 5’-TTCTCTTCTGCGTGAATGCACCAGATGA-3’TAMRA). Each 20 μl reaction contained 900 nM of each primer and 250 nM of probe, and 1x Fast Virus 1-Step Master Mix, plasma-derived vRNA sample, SIV gag RNA transcript containing standard, or no template control.

qRT-PCR was performed in a ABI Step One Plus Cycler. PCR was performed with an initial step at 50°C for 5 min followed by a second step at 95°C for 20 sec, and then 40 cycles of 95°C for 15 sec and 60°C for 1 min. Ten-fold serial dilutions (1 to 1 × 10^6^ copies per reaction) of an in vitro transcribed SIV gag RNA were used to generate standard curves. Each sample was tested in duplicate reactions. Plasma viral loads were calculated and shown as viral RNA copies/mL plasma. The limit of detection is 50 copies/ml. Infections were confirmed after a consecutive positive plasma viral load measurement.

The following tissues were biopsied: inguinal lymph nodes, rectum and vagina. Tissues were snap-frozen and stored at -80 °C until further analysis of cell-associated SHIV DNA. To detect viral DNA in tissue samples, total DNA was isolated from PBMCs or tissue specimens using the Qiagen DNeasy Blood & Tissue Kit (Qiagen #69504) according to the manufacturer’s protocol. DNA was quantified by RNAseP qRT-PCR assay according to the manufacturer’s protocol using TaqMan Copy Number Reference Assay (ThermoFisher #4403326), TaqMan Genotyping Master Mix (ThermoFisher #4371357), and serial 10-fold dilutions of Taqman Control Genomic DNA (ThermoFisher#4312660) for a standard curve. To quantify cell-associated SHIV DNA, qRT-PCR was performed using the SIV gag primer probe set described above. Each 20 μl reaction contained 900 nM of each primer and 250 nM of probe, and 1x TaqMan Gene Expression Master Mix (Applied Biosystems, Foster City, CA), macaque-derived DNA sample, SIV gag DNA containing standard, or no template control. PCR was initiated in with an initial step of 50°C for 2 min and then 95°C for 10 min. This was followed by 40 cycles of 95°C for 15 sec, and 60°C for 1 min. Each sample was tested in triplicate reactions. Ten-fold serial dilutions of a SIV gag DNA template (1 to 1 × 10^5^ per reaction) were used to generate standard curves. The limit of detection of this assay was determined to be 1 copy of SIV gag DNA.

### Device retrieval

Implants (nISL and PBS) were retrieved on day 112 from the PrEP-treated macaques (n=11) for continuation to a 2-month drug-washout period. SHIV-infected macaques in the control group (n=12) were transferred to another study (data not shown) and euthanized 28 days later. The implant with surrounding fibrotic capsule was retrieved with a small incision in the skin and was sutured with a simple interrupted tacking suture of 4-0 PDS. The implant was removed from within the fibrotic capsule via a small incision. A 3D printed implant made with flexible polymer and identical dimensions was placed within the fibrotic capsule as a placeholder for tissue fixation. The tissue was fixed in 10% buffered formalin and stored in 70% ethanol. The tissue was sectioned, and the placeholder implant removed prior to histological analysis. Tissues surrounding the implant site were then embedded in paraffin, cut into 5 μm sections and stained with H&E and Masson Trichrome staining at the Research Pathology Core HMRI, Houston, TX, USA.

### Residual drug quantification from explanted implants

Upon explantation, the implants were stored at -80 °C until further analysis. For residual drug retrieval, the implants were thawed at 4°C overnight. A hole was drilled on the outermost corner on the back of the implant using a 3/64 titanium drill bit with a stopper. Drilling was performed on the membrane shell under the port with no titanium filter. Following drilling, the implants were placed in 50 mL conical tubes with 20 mL of 70% ethanol. Each implant was flushed using a 19-gauge needle with 70% ethanol from the sink solution. For sterilization, the implants were incubated in 70% ethanol for 4 days. Next, since ISL did not solubilize, ethanol was evaporated under vacuum and 20 mL of dimethylsulfoxide (DMSO) were added to each tube. Tubes were left shaking overnight. The DMSO samples were transferred to 0.2 μm nylon centrifugal filters and centrifuged at 500 G for 8 minutes at room temperature. An aliquot of 10 μL from the filtered samples were further diluted in 990 μL 1X PBS. High-performance liquid chromatography (HPLC) analysis was performed on an Agilent Infinity 1260 system equipped with a diode array and evaporative light scattering detectors using a 3.5-μm 4.6 × 100 mm Eclipse Plus C18 column and water/methanol as the eluent and 25 μL injection volume, as previously described. Specifically, ammonia acetate buffered water (solvent A) and ammonia acetate buffered methanol (solvent B) at 2.00 mL/min flow rate gradient: 0 min (5% B), 0.8 min (5% B), 3.8 min (100% B), 4.6 min (100% B), 5.2 min (5% B). Peak areas were analyzed at 261 nm absorbance.

### Assessment of PrEP nISL safety and tolerability

Semiquantitative histopathological assessment of inflammatory response to a foreign body was evaluated in accordance to the inflammation scoring system presented in Su *et al*.(*21*), which was adopted from a published standard. Briefly, cells were counted via high power field (HPF) and scored (0-4) based on histological characteristics: polymorphonuclear cells, lymphocytes, plasma cells, macrophages, giant cells, necrosis, capsule thickness, and tissue infiltrate. The scores reported by each pathologist were averaged per implant (see Tables S2, S3, Supplementary Materials). Then, the total histological characteristic scores were reported per group as the average of the sum of all histological scores of all implants. The reactivity grade for each implant was computed using Equation 1 from Su *et al*.(*21*) and the average placebo-adjusted implant reactivity score (S_pair_) was calculated by subtracting the result obtained for PBS from nISL. The S_pair_ classification used in Su *et al*.(*21*) and published standard was adopted: minimal to no reaction (0.0 < S_pair_ < 2.9), slight reaction (3.0 < S_pair_ < 8.9), moderate reaction (9.0 < S_pair_ < 15.0), and severe reaction (S_pair_ > 15.1).

To assess ISL toxicity, a comprehensive metabolic panel and CBC was analyzed for each animal monthly during all studies to assess kidney and liver function and monitor the well-being of the NHPs.

### FRAP analysis in fibrotic capsules

The tissue diffusion coefficient measurements were performed according to our previous publication. Briefly, formalin-fixed fibrotic capsule tissue samples (n=2 per group) and subcutaneous tissue samples (n=2) were placed in a solution of 2 mg/ml FITC (Invitrogen) in DMEM/F12 media (Gibco). After overnight incubation at 4°C, the samples were removed from the solution and placed in microscope chamber slides (Ibidi). FRAP (fluorescence recovery after photobleaching) measurement were performed with the Olympus Fluoview 3000 confocal microscope equipped with a 10x objective and a 488 nm laser on 3 FOV per sample. Measurements of the FITC solution alone (n=3) were performed to validate the method. For each FOV, three pre-bleaching images were collected at low laser intensity (1%). Then, bleaching was achieved by exposing a circular area (100 pixels or 248 μm diameter) at 100% laser intensity for 60 seconds. Finally, fluorescence recovery was assessed by acquiring images of the bleached area at low laser intensity for 15 minutes (10 images/minute).

The data was analyzed with the Olympus cellSens software to obtain the characteristic diffusion time τ. This value, along with the radius of the bleached area, ω, is used to calculate the diffusion coefficient, D, according to the following equation(*86*):

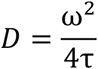

### Imaging Mass Cytometry (IMC)

IMC analysis was conducted at the ImmunoMonitoring Core at the Houston Methodist Research Institute. Antibodies were conjugated with metals following the Fluidigm protocol as indicated elsewhere (*87*) (Supplementary Material Table S4). For staining, tissue sections were kept at 60 °C overnight followed by deparaffinization in xylene and rehydration in alcohol gradients (absolute ethanol, absolute ethanol : deionized water 90:10, 80:20, 70:30, 50:50, 0:100) for 10 min each. Epitope retrieval was performed in a water bath at 95 °C in Tris-Tween 20 buffer pH 9 for 20 min. Sections were blocked with 3% BSA in tris-buffered saline, followed by staining overnight at 4 °C with markers in Supplementary Material Table S4 and nuclear staining with Cell-ID Intercalator (Fluidigm). Specifically, 22 markers and marker clusters were used to map immune cell populations. The panel included cell surface receptors (CD45, CD4, CD8, CD20), signaling mediators (STAT1, STAT3), markers of cell activation (NFkB), vascular cell adhesion (VCAM1), differentiation of M1 to M2 cells (CD68+CD163+) or indicators of cell proliferation (Ki67). Clusters characterized by co-expression of myeloperoxidase (MPO) and Granzyme B (GranZB), HLADR+CD2-+Ki67+ or co-expression of phenotype markers CD3, CD4 or CD8 combined with markers for cell proliferation (Ki67), activation (tBet), or indicators of T cell regulatory populations (Foxp3), activated CD8 cells (GranZB+) or phenotypic markers that distinguish effector T cells from exhausted T cells (PD1+) were also analyzed. After staining, slides were air-dried and ablated with Hyperion (Fluidigm) for data acquisition. Data were segmented by ilastik and CellProfiler. Histology topography cytometry analysis toolbox (HistoCAT) and R scripts were used to quantify cell number and generate tSNE plots(*88*). Analysis was performed on 2 ROI/sample and *n* = 2/group for a total of *n* = 3 ROI/group.

### Quantification of inflammatory cytokines from fluid collected from implants

After explantation, fluid was extracted from the drug reservoir using a syringe with 21G needle. Levels of inflammatory cytokines were quantified using Bio-Plex Pro Human Inflammation Panel 1 37-plex (#171AL001M) magnetic bead-based assay (Bio-Rad Laboratories) according to manufacturer’s instructions. Briefly, fluid samples were centrifuged for 10 min at 10,000g for clarification. Supernatant was diluted (1:4) with sample diluent HB to a final concentration of 0.5% BSA. 50 μl of 1x magnetic beads suspension was added to each well, followed by 50 μl of diluted sample. Each sample analysis was performed in replicates (n=2 or 3) depending on sample volume availability. Cytokine levels were assayed on the Bio-Plex® 200 System (Bio-Rad). Cytokine concentrations were calculated using an 8-point calibration curve for each individual analyte produced from manufacturer-supplied set standards of known concentration (pg/ml).

### Statistical analysis

Data are represented as mean ± SD or median with interquartile range (IQR) between the first (25th percentile) and third (75th percentile) quartiles. The Kaplan-Meier analysis was performed between the PrEP and control groups, with the use of the number of inoculations as the time variable. The exact log-rank test was used to test the survival between the two groups. Unpaired t test and one-way ANOVA were used to compare between groups. Outliers were identified via ROUT method with Q=0.1%. Statistical significance was defined as two-tailed p<0.05 for all tests. All statistical analyses were performed with GraphPad Prism 9 (version 9.3.1; GraphPad Software, Inc., La Jolla, CA).

## List of Supplementary Materials

Fig. S1 to Fig. S11

Table S1 to Table S4

## Acknowledgements

We thank Dr. Andreana L. Rivera, Yuelan Ren, and Sandra Steptoe from the research pathology core of Houston Methodist Research Institute. Dr. Jianhua “James” Gu from the electron microscopy core. We thank Luke Segura, Dana Salazar, Elizabeth Lindemann and Dr. Greg Wilkerson from the Michael E. Keeling Center for Comparative medicine and Research at UTMDACC for support in animal studies. We thank Drs. Gerardo Garcia-Lerma and Charles Dobard at the Centers for Disease Control and Prevention for detailed discussions on viral-challenge analysis in NHP models.

## Funding

This work was supported by funding from:

National Institutes of Health National Institute of Allergy and Infectious Diseases (R01AI120749; A.G.)

National Institutes of Health National Institute of General Medical Sciences (R01GM127558; A.G.)

Basic Science Core of the Texas Developmental Center for AIDS Research (P30 AI161943; J.T.K.).

## Author contributions

Conceptualization: FPPF, RCA, AG

Methodology: FPPF, NDT, LRB, JTK, PLA

Investigation: FPPF, NDT, BN, SS, KS, LRB, FA, MW, LR, DN, MMI, JTK, PNN, AG

Visualization: FPPF, NDT, AG

Funding acquisition: AG

Project administration: FPPF, PNN, AG

Supervision: AG

Validation: LRB, JTK, PLA, AG

Resources: JTK, PLA, PNN, AG

Writing – original draft: FPPF, NDT, CYXC, AG

Writing – review & editing: FPPF, NDT, CYXC, JTK, PLA, RCA, AG

## Competing interests

A.G. is an inventor of intellectual property licensed by Semper Therapeutics. A.G. and P.L.A. receives grants and contracts from Gilead Sciences paid to their respective institutions. P.L.A. collects personal fees from Gilead Sciences. All other authors declare that they have no competing interests.

## Data and materials availability

All data are available in the main text or the supplementary materials.

## Correspondence

should be addressed to A.G.

