## Supplementary Materials for "Ultra-long-acting refillable nanofluidic implant confers full protection against SHIV infection in non-human primates"

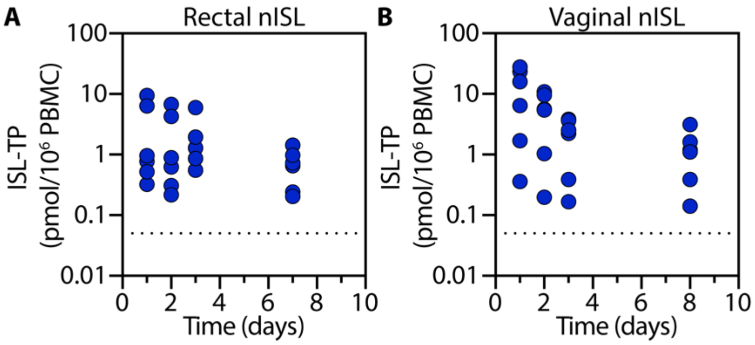

Fig. S1. ISL-TP PBMC concentrations (pmol/$10^6$ PBMC) in rectal and vaginal efficacy study surpass preventive threshold by day 1. Preventive threshold of 0.05 pmol/106 PBMC (horizontal dashed line).

### 1. Implant safety and tolerability

Of note, for the implant that flipped during the long-term PK study, we observed a significantly thicker fibrotic capsule as compared to control PBS implants or nISL implants releasing ISL toward the subcutaneous tissues (Fig. S2, A, B). The fibrotic capsule showed massive influx of immune cells (Fig. S2 D-G). Specifically, lymphocytes and macrophages were observed (Fig. S2 D, F), which are indicative of an on-going chronic inflammation. In addition, localized necrosis (Fig. S2 C), activated myofibroblasts and presence of giant cells (Fig. S2 E, G), were visible providing additional evidence of the severe FBR. Overall, the substantial inflammatory response in this sample aligns with the visual observation of local swelling at the implant site. Importantly, this immune analysis supports our hypothesis that ISL could induce tissue inflammation when continuously released towards the skin, where there is less vascularity and thus reduced drug transport.

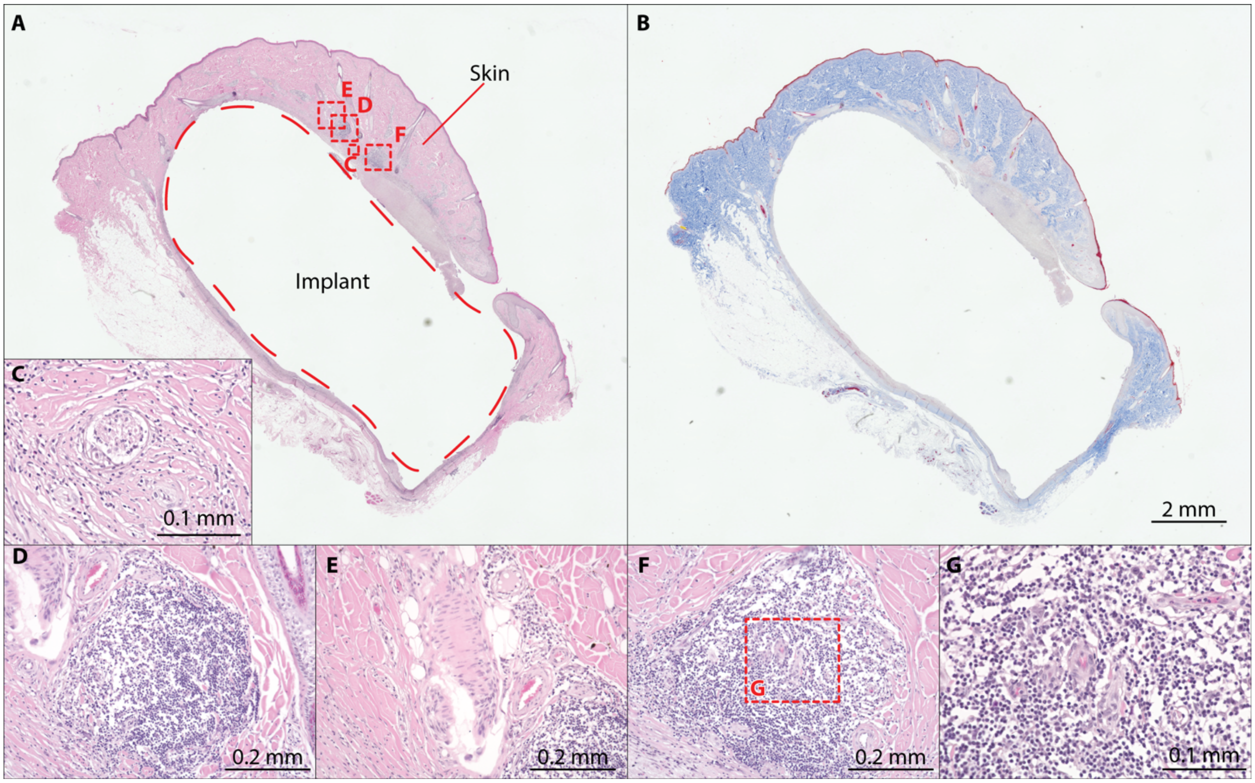

Fig S2. Fibrotic capsule histology of implant that flipped in the long term PK. (**A**) H&E and (**B**) Masson trichrome staining. (**C**) 40X magnification of area indicated in A. (**D-F**) 20X magnification of area indicated in A. (**G**) 40X magnification of area indicated in F.

| Adverse Events | Control PBS n=12 (%) | ISL rectal n=6 (%) | ISL vaginal n=6 (%) |
|---|---|---|---|
| **General disorders** | | | |
| Implant site erythema | | | |
| Implant site induration | | 5 (83%) | 3 (50%) |
| Implant site pruritus | | | |
| Implant site hematoma | | | |
| **Procedure complication** | | | |
| Wound complication | | | |
| Wound secretion | | | |

Table S1. NHP clinical observations tolerability to ISL-releasing implants.

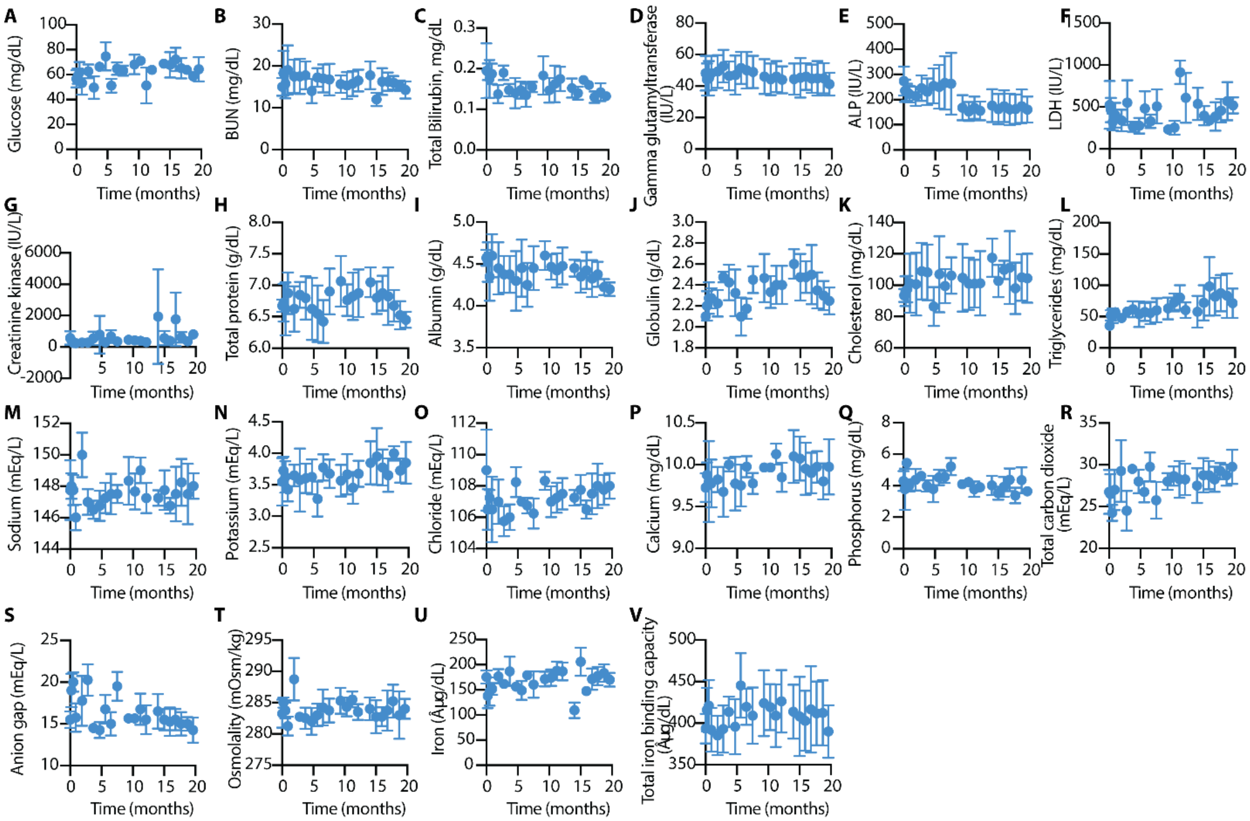

Fig. S3. (A-V) Metabolic panel of rhesus macaques with nISL in long-term PK study. Baseline value for comparison is on day 0 pre-implantation. All data presented as mean ± SD.

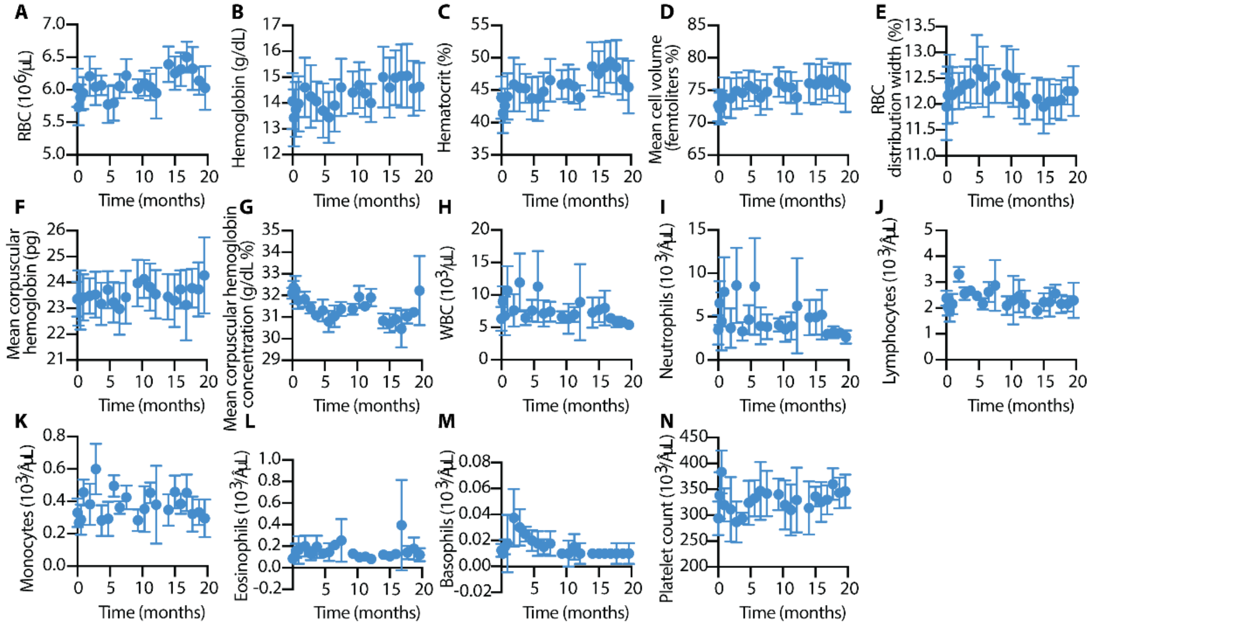

Fig. S4. (A-N) CBC of rhesus macaques with nISL in PK study. Baseline value for comparison is on day 0 pre-implantation. All data presented as mean ± SD.

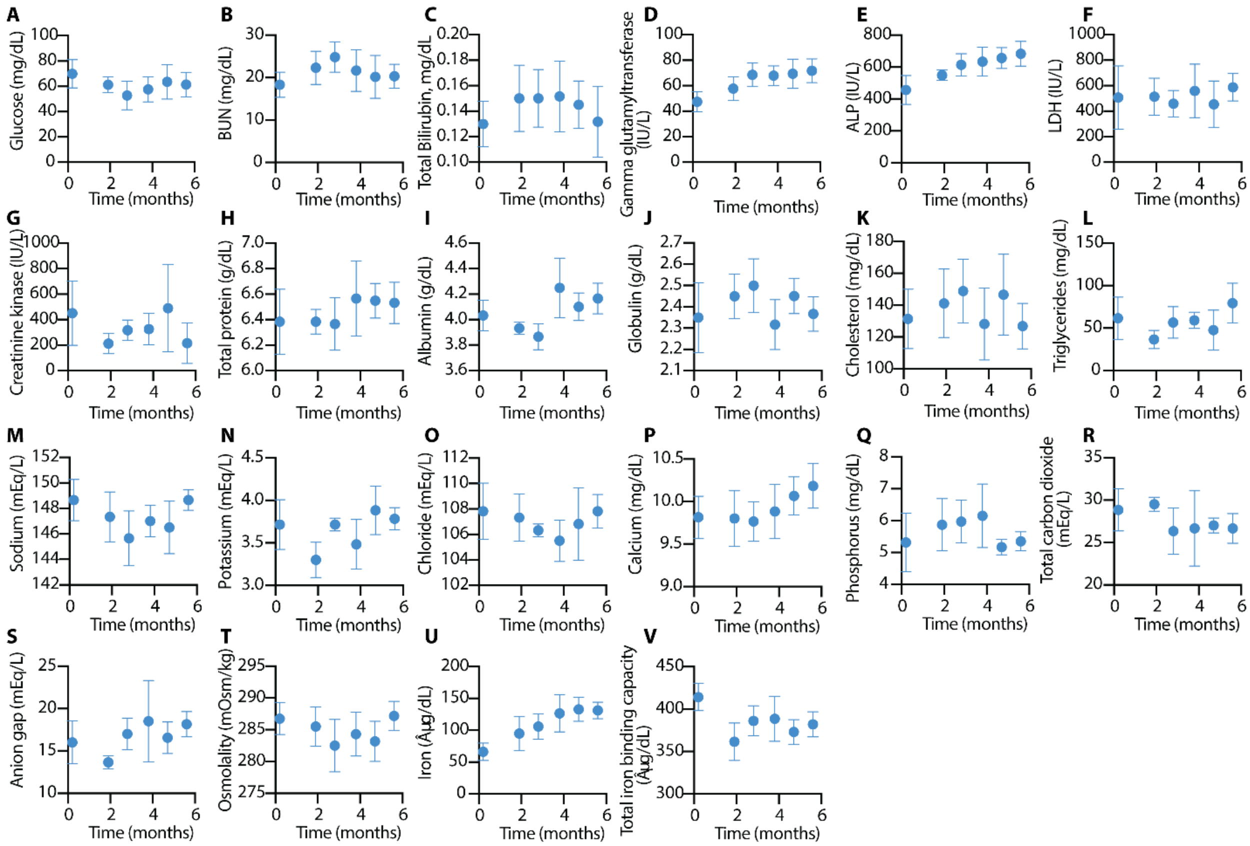

Fig. S5. (A-V) Metabolic panel of rhesus macaques with nISL in rectal efficacy study. Baseline value for comparison is on day 0 pre-implantation. All data presented as mean ± SD.

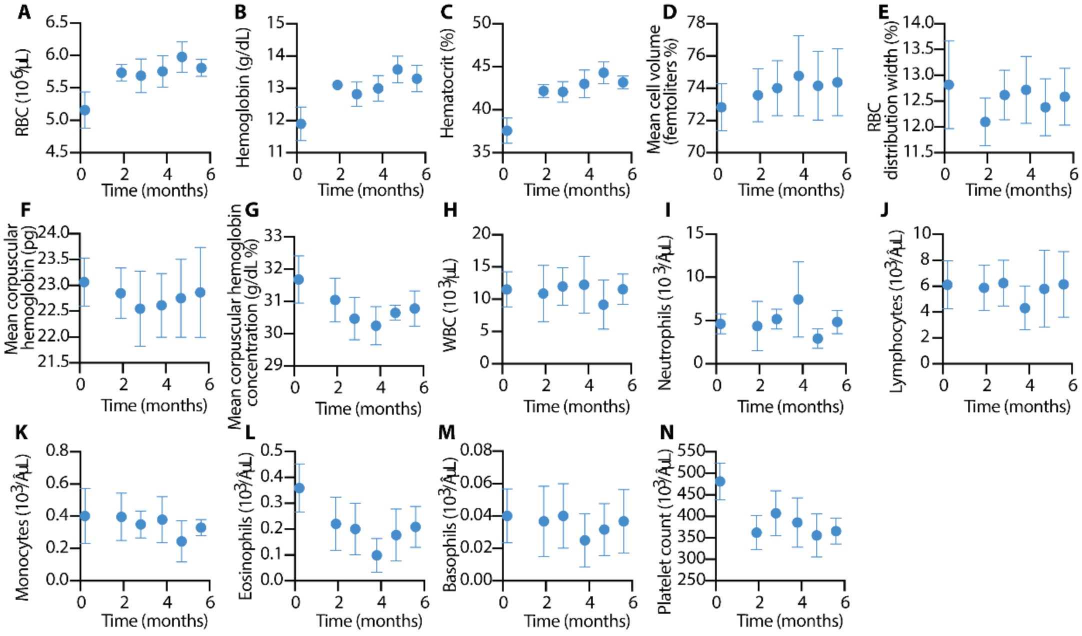

Fig. S6. (A-N) CBC of rhesus macaques with nISL in rectal efficacy study. Baseline value for comparison is on day 0 pre-implantation. All data presented as mean ± SD.

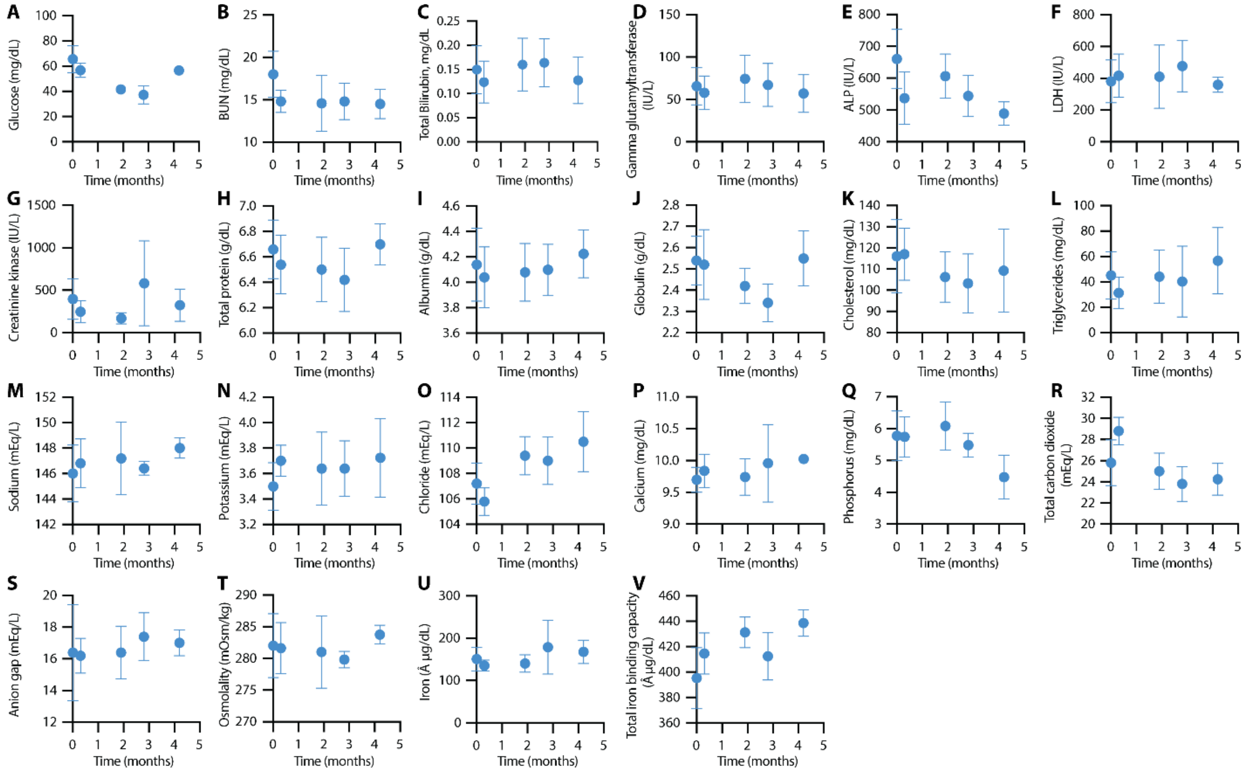

Fig. S7. (A-V) Metabolic panel of rhesus macaques with nISL in vaginal efficacy study. Baseline value for comparison is on day 0 pre-implantation. All data presented as mean ± SD.

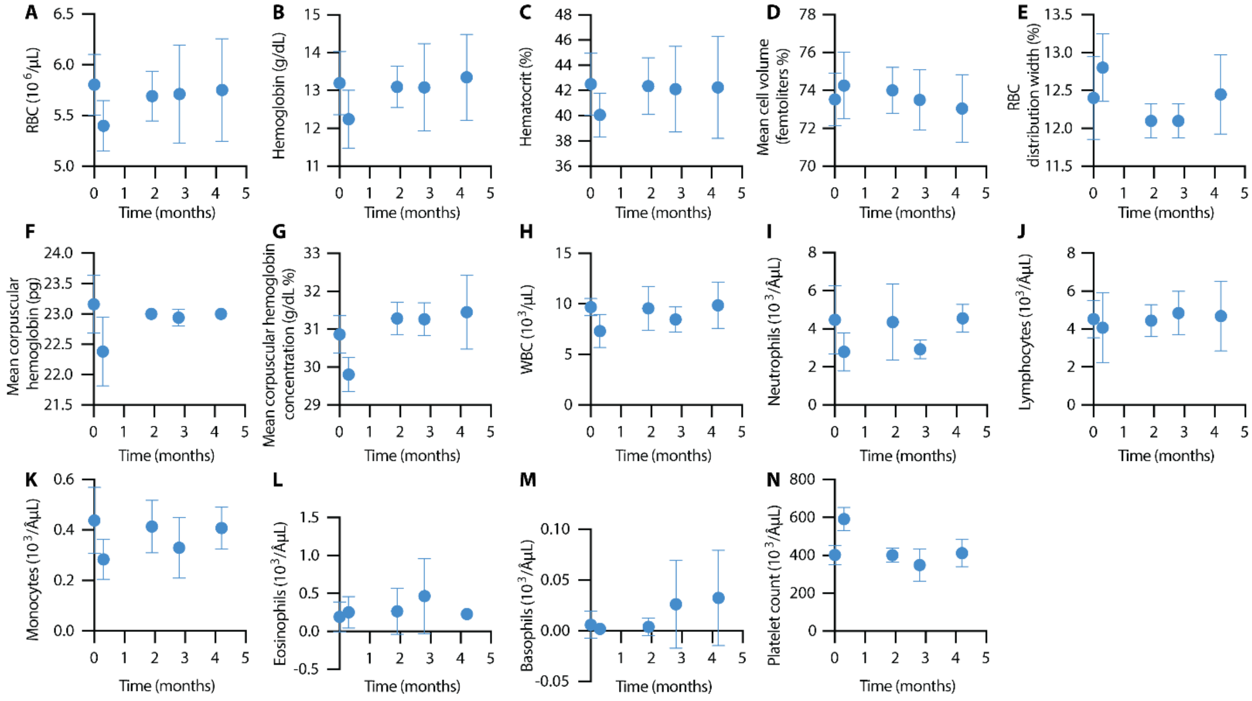

Fig. S8. (A-N) CBC of rhesus macaques with nISL vaginal efficacy study. Baseline value for comparison is on day 0 pre-implantation. All data presented as mean ± SD.

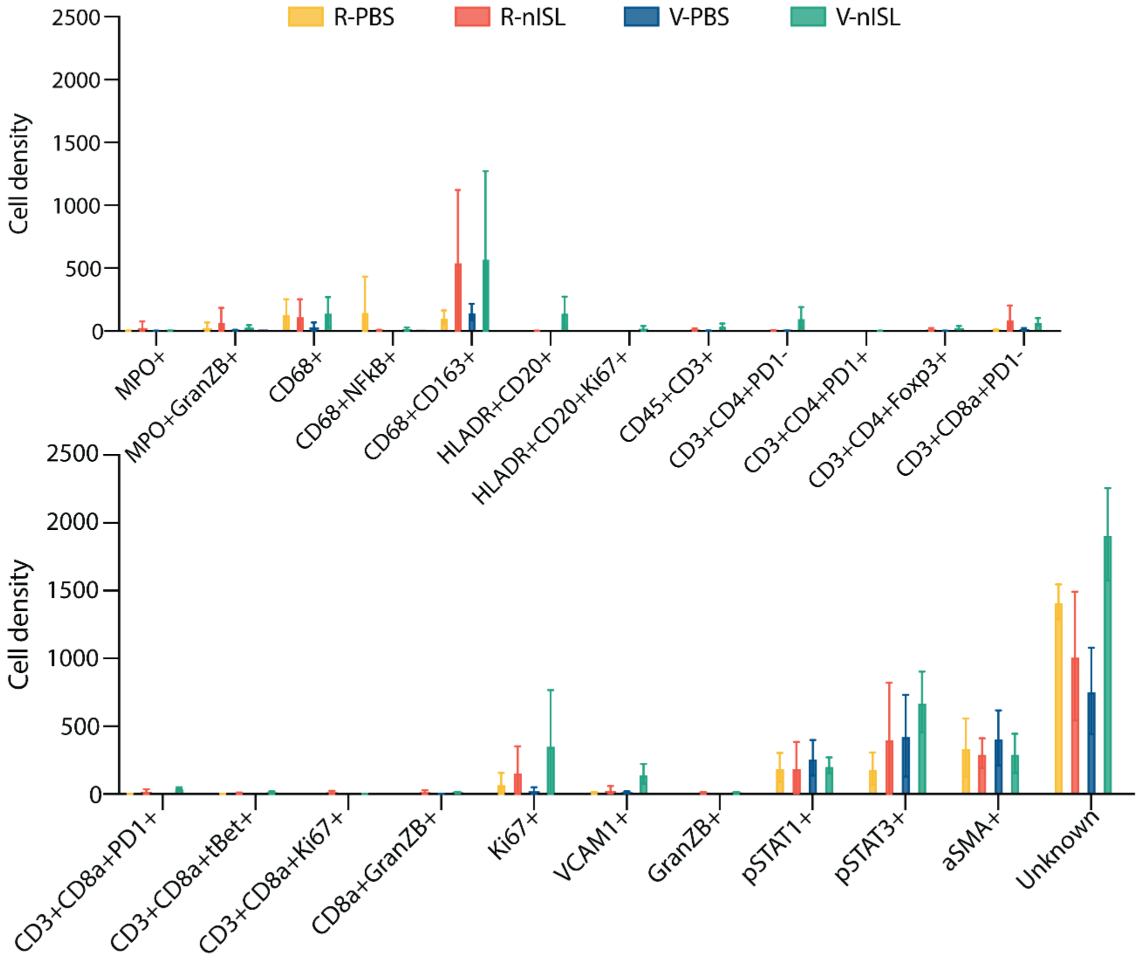

Fig S9. IMC cell densities for R-PBS, R-nISL, V-PBS and V-nISL.

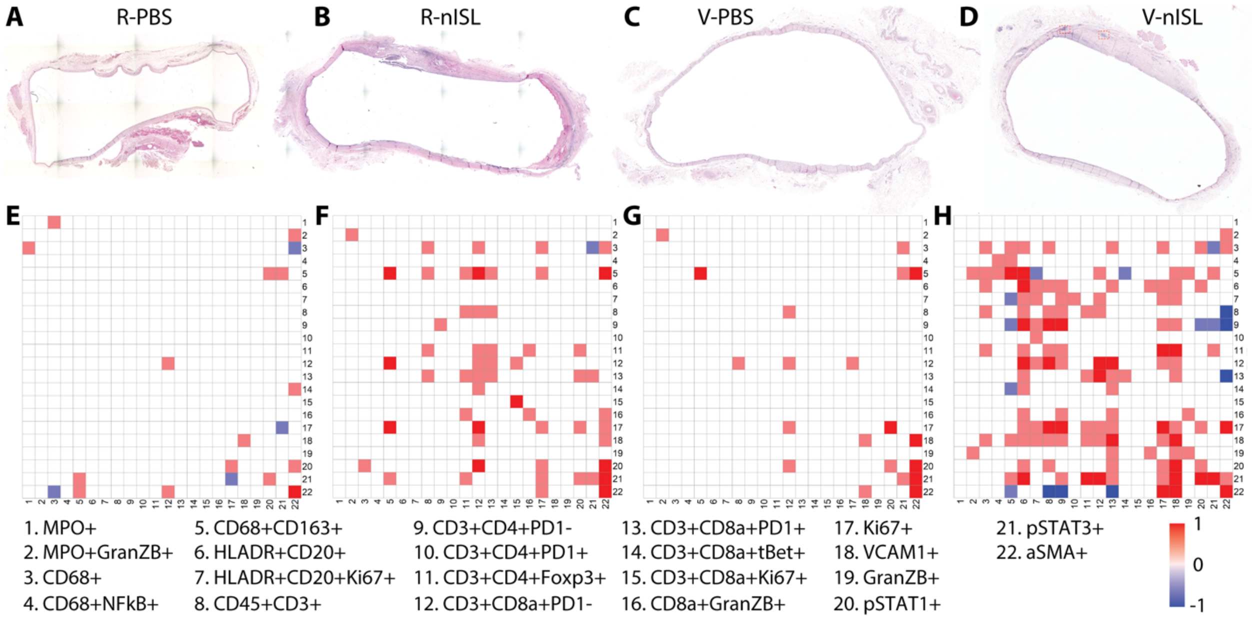

Fig S10. H&E staining of entire fibrotic capsule tissue. (**A**) R-PBS, (**B**) R-nISL, (**C**) V-PBS, (**D**) V-nISL. Neighborhood analysis of IMC data for representative sample of (**E**) R-PBS, (**F**) R-nISL, (**G**) V-PBS, (**H**) V-nISL.

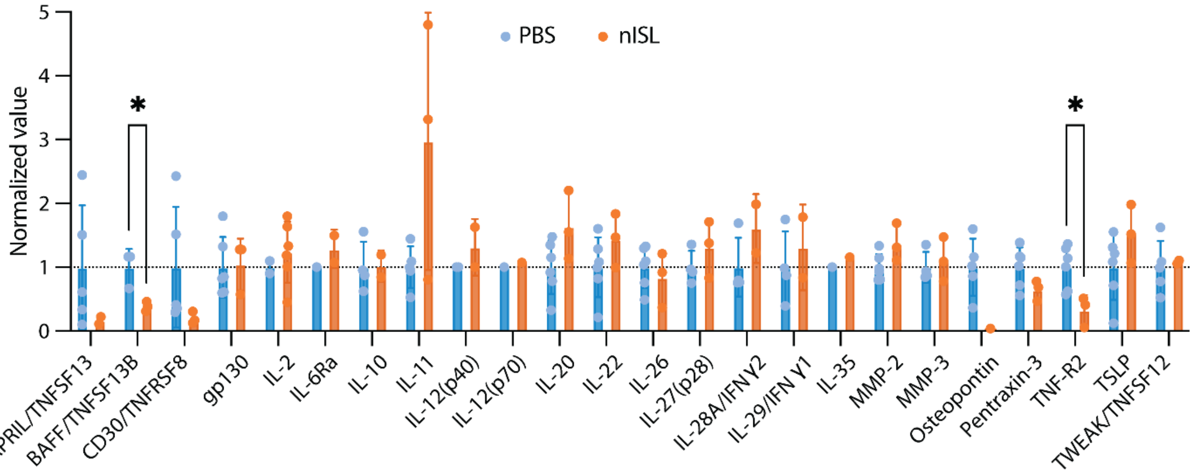

Fig S11. Bioplex of interstitial fluid collected from explanted devices. All data presented as mean ± SD.

| **19-223 R-PBS** | **Score** | | |
|---|---|---|---|
| | **Pathologist 1** | **Pathologist 2** | **Average** |
| Polymorphonuclear cells | 3 | 2 | **2.5** |
| Lymphocytes | 1 | 1 | **1.0** |
| Plasma cells | 1 | 0 | **0.5** |
| Macrophages | 1 | 1 | **1.0** |

| | | | |
|---|---|---|---|
| Giant cells | 0 | 0 | **0.0** |
| Necrosis | 0 | 0 | **0.0** |
| Capsule thickness | 2.5 | 2 | **2.3** |
| Tissue infiltrate | 1 | 1 | **1.0** |
| **Overall total** | **9.5** | **7** | **8.3** |
| **19-223 R-nISL** | **Score** | | |
| | **Pathologist 1** | **Pathologist 2** | **Average** |
| Polymorphonuclear cells | 0 | 0 | **0.0** |
| Lymphocytes | 2 | 1 | **1.5** |
| Plasma cells | 0 | 0 | **0.0** |
| Macrophages | 1 | 0 | **0.5** |
| Giant cells | 0 | 0 | **0.0** |
| Necrosis | 0 | 2 | **1.0** |
| Capsule thickness | 3 | 4 | **3.5** |
| Tissue infiltrate | 1 | 0 | **0.5** |
| **Overall total** | **7** | **7** | **7.0** |
| **19-100 R-PBS** | **Score** | | |
| | **Pathologist 1** | **Pathologist 2** | **Average** |
| Polymorphonuclear cells | 0 | 0 | **0.0** |
| Lymphocytes | 1 | 1 | **1.0** |
| Plasma cells | 0 | 0 | **0.0** |
| Macrophages | 0 | 0 | **0.0** |
| Giant cells | 0 | 0 | **0.0** |
| Necrosis | 0 | 0 | **0.0** |
| Capsule thickness | 2 | 2 | **2.0** |
| Tissue infiltrate | 0 | 0 | **0.0** |
| **Overall total** | **3** | **3** | **3.0** |
| **19-100 R-nISL** | **Score** | | |
| | **Pathologist 1** | **Pathologist 2** | **Average** |
| Polymorphonuclear cells | 0 | 0 | **0.0** |
| Lymphocytes | 3 | 1 | **2.0** |
| Plasma cells | 1 | 0 | **0.5** |
| Macrophages | 1 | 1 | **1.0** |
| Giant cells | 0 | 0 | **0.0** |
| Necrosis | 0 | 1 | **0.5** |
| Capsule thickness | 2 | 2 | **2.0** |

| | | | |
|---|---|---|---|
| Tissue infiltrate | 3 | 2 | **2.5** |
| **Overall total** | **10** | **7** | **8.5** |
| **19-226 R-PBS** | **Score** | | |
| | **Pathologist 1** | **Pathologist 2** | **Average** |
| Polymorphonuclear cells | 0 | 0 | **0.0** |
| Lymphocytes | 1 | 0 | **0.5** |
| Plasma cells | 0 | 0 | **0.0** |
| Macrophages | 0 | 0 | **0.0** |
| Giant cells | 0 | 0 | **0.0** |
| Necrosis | 0 | 0 | **0.0** |
| Capsule thickness | 2 | 1 | **1.5** |
| Tissue infiltrate | 0 | 0 | **0.0** |
| **Overall total** | **3** | **1** | **2.0** |
| **19-226 R-nISL** | **Score** | | |
| | **Pathologist 1** | **Pathologist 2** | **Average** |
| Polymorphonuclear cells | 2 | 2 | **2** |
| Lymphocytes | 2 | 1 | **1.5** |
| Plasma cells | 0 | 0 | **0** |
| Macrophages | 2 | 1 | **1.5** |
| Giant cells | 1 | 0 | **0.5** |
| Necrosis | 2 | 2 | **2** |
| Capsule thickness | 3 | 3 | **3** |
| Tissue infiltrate | 1 | 2 | **1.5** |
| **Overall total** | **13** | **11** | **12** |
| **19-225 R-PSB** | **Score** | | |
| | **Pathologist 1** | **Pathologist 2** | **Average** |
| Polymorphonuclear cells | 0 | 0 | **0.0** |
| Lymphocytes | 2 | 1 | **1.5** |
| Plasma cells | 0 | 0 | **0.0** |
| Macrophages | 2 | 2 | **2.0** |
| Giant cells | 1 | 1 | **1.0** |
| Necrosis | 0 | 0 | **0.0** |
| Capsule thickness | 2 | 2 | **2.0** |
| Tissue infiltrate | 0 | 0 | **0.0** |
| **Overall total** | **7** | **6** | **6.5** |

| 19-225 R-nISL | Score | | |
|---|---|---|---|
| | **Pathologist 1** | **Pathologist 2** | **Average** |
| Polymorphonuclear cells | 2 | 2 | **2.0** |
| Lymphocytes | 2 | 2 | **2.0** |
| Plasma cells | 0 | 0 | **0.0** |
| Macrophages | 1 | 2 | **1.5** |
| Giant cells | 0 | 0 | **0.0** |
| Necrosis | 1 | 1 | **1.0** |
| Capsule thickness | 2 | 3 | **2.5** |
| Tissue infiltrate | 1 | 1 | **1.0** |
| **Overall total** | **9** | **11** | **10.0** |
| **19-238 R-PBS** | **Score** | | |
| | **Pathologist 1** | **Pathologist 2** | **Average** |
| Polymorphonuclear cells | 0 | 0 | **0.0** |
| Lymphocytes | 1 | 0 | **0.5** |
| Plasma cells | 0 | 0 | **0.0** |
| Macrophages | 1 | 2 | **1.5** |
| Giant cells | 0 | 0 | **0.0** |
| Necrosis | 0 | 0 | **0.0** |
| Capsule thickness | 1 | 1 | **1.0** |
| Tissue infiltrate | 0 | 0 | **0.0** |
| **Overall total** | **3** | **3** | **3.0** |
| **19-238 R-nISL** | **Score** | | |
| | **Pathologist 1** | **Pathologist 2** | **Average** |
| Polymorphonuclear cells | 2 | 2 | **2.0** |
| Lymphocytes | 2 | 2 | **2.0** |
| Plasma cells | 0 | 0 | **0.0** |
| Macrophages | 1 | 2 | **1.5** |
| Giant cells | 0 | 0 | **0.0** |
| Necrosis | 1 | 2 | **1.5** |
| Capsule thickness | 2 | 3 | **2.5** |
| Tissue infiltrate | 1 | 0 | **0.5** |
| **Overall total** | **9** | **11** | **10.0** |
| **19-231 R-PBS** | **Score** | | |
| | **Pathologist 1** | **Pathologist 2** | **Average** |
| Polymorphonuclear cells | 1 | 0 | **0.5** |

| | | | |
|---|---|---|---|
| Lymphocytes | 1 | 0 | **0.5** |
| Plasma cells | 0 | 0 | **0.0** |
| Macrophages | 1 | 2 | **1.5** |
| Giant cells | 0 | 0 | **0.0** |
| Necrosis | 0 | 0 | **0.0** |
| Capsule thickness | 2 | 2 | **2.0** |
| Tissue infiltrate | 0 | 0 | **0.0** |
| **Overall total** | **5** | **4** | **4.5** |
| **19-231 R-nISL** | **Score** | | |
| | **Pathologist 1** | **Pathologist 2** | **Average** |
| Polymorphonuclear cells | 2.5 | 1 | **1.8** |
| Lymphocytes | 2.5 | 2 | **2.3** |
| Plasma cells | 0 | 0 | **0.0** |
| Macrophages | 2 | 2 | **2.0** |
| Giant cells | 0 | 0 | **0.0** |
| Necrosis | 3 | 3 | **3.0** |
| Capsule thickness | 3 | 2 | **2.5** |
| Tissue infiltrate | 2 | 2 | **2.0** |
| **Overall total** | **15** | **12** | **13.5** |

Table S2. Histopathological scores of R-nISL (PrEP) and R-PBS (control) implants. All pathologists are board-certified. Pathologist 1and 2 are from Houston Methodist Hospital and Baylor College of Medicine, respectively.

| **19-037 V-PBS** | **Score** | | |
|---|---|---|---|
| | **Pathologist 1** | **Pathologist 2** | **Average** |
| Polymorphonuclear cells | 0 | 0 | **0.0** |
| Lymphocytes | 2 | 2 | **2.0** |
| Plasma cells | 1 | 2 | **1.5** |
| Macrophages | 2 | 2 | **2.0** |
| Giant cells | 0 | 0 | **0.0** |
| Necrosis | 0 | 1 | **0.5** |
| Capsule thickness | 3 | 2 | **2.5** |
| Tissue infiltrate | 2 | 2 | **2.0** |
| **Overall total** | **10** | **11** | **10.5** |
| **19-046 V-PBS** | **Score** | | |
| | **Pathologist 1** | **Pathologist 2** | **Average** |
| Polymorphonuclear cells | 0 | 0 | **0.0** |

| | | | |
|---|---|---|---|
| Lymphocytes | 1 | 1 | **1.0** |
| Plasma cells | 0 | 0 | **0.0** |
| Macrophages | 1 | 1 | **1.0** |
| Giant cells | 0 | 0 | **0.0** |
| Necrosis | 0 | 0 | **0.0** |
| Capsule thickness | 2 | 1 | **1.5** |
| Tissue infiltrate | 0 | 0 | **0.0** |
| **Overall total** | **4** | **3** | **3.5** |
| **19-054 V-PBS** | **Score** | | |
| | **Pathologist 1** | **Pathologist 2** | **Average** |
| Polymorphonuclear cells | 0 | 0 | **0.0** |
| Lymphocytes | 2 | 1 | **1.5** |
| Plasma cells | 0 | 0 | **0.0** |
| Macrophages | 1 | 0 | **0.5** |
| Giant cells | 0 | 0 | **0.0** |
| Necrosis | 0 | 0 | **0.0** |
| Capsule thickness | 2 | 1 | **1.5** |
| Tissue infiltrate | 1 | 0 | **0.5** |
| **Overall total** | **6** | **2** | **4.0** |
| **19-068 V-PBS** | **Score** | | |
| | **Pathologist 1** | **Pathologist 2** | **Average** |
| Polymorphonuclear cells | 0 | 0 | **0.0** |
| Lymphocytes | 1 | 0 | **0.5** |
| Plasma cells | 0 | 0 | **0.0** |
| Macrophages | 1 | 0 | **0.5** |
| Giant cells | 0 | 0 | **0.0** |
| Necrosis | 0 | 0 | **0.0** |
| Capsule thickness | 1 | 1 | **1.0** |
| Tissue infiltrate | 0 | 0 | **0.0** |
| **Overall total** | **3** | **1** | **2.0** |
| **19-074 V-PBS** | **Score** | | |
| | **Pathologist 1** | **Pathologist 2** | **Average** |
| Polymorphonuclear cells | 0 | 0 | **0.0** |
| Lymphocytes | 2 | 0 | **1.0** |
| Plasma cells | 0 | 0 | **0.0** |
| Macrophages | 1 | 1 | **1.0** |

| | | | |
|---|---|---|---|
| Giant cells | 0 | 0 | **0.0** |
| Necrosis | 0 | 0 | **0.0** |
| Capsule thickness | 1 | 1 | **1.0** |
| Tissue infiltrate | 1 | 0 | **0.5** |
| **Overall total** | **5** | **2** | **3.5** |
| **19-087 V-PBS** | **Score** | | |
| | **Pathologist 1** | **Pathologist 2** | **Average** |
| Polymorphonuclear cells | 1 | 0 | **0.5** |
| Lymphocytes | 2 | 1 | **1.5** |
| Plasma cells | 1 | 0 | **0.5** |
| Macrophages | 2 | 2 | **2** |
| Giant cells | 0 | 0 | **0** |
| Necrosis | 0 | 1 | **0.5** |
| Capsule thickness | 2 | 1 | **1.5** |
| Tissue infiltrate | 1 | 2 | **1.5** |
| **Overall total** | **9** | **7** | **8** |
| **19-037 V-nISL** | **Score** | | |
| | **Pathologist 1** | **Pathologist 2** | **Average** |
| Polymorphonuclear cells | 1 | 0 | **0.5** |
| Lymphocytes | 3 | 2 | **2.5** |
| Plasma cells | 1 | 1 | **1.0** |
| Macrophages | 3 | 3 | **3.0** |
| Giant cells | 0 | 0 | **0.0** |
| Necrosis | 1 | 1 | **1.0** |
| Capsule thickness | 4 | 4 | **4.0** |
| Tissue infiltrate | 2 | 3 | **2.5** |
| **Overall total** | **15** | **14** | **14.5** |
| **19-046 V-nISL** | **Score** | | |
| | **Pathologist 1** | **Pathologist 2** | **Average** |
| Polymorphonuclear cells | 1 | 0 | **0.5** |
| Lymphocytes | 2 | 1 | **1.5** |
| Plasma cells | 0 | 0 | **0.0** |
| Macrophages | 2 | 2 | **2.0** |
| Giant cells | 0 | 0 | **0.0** |
| Necrosis | 0 | 0 | **0.0** |
| Capsule thickness | 3 | 4 | **3.5** |

| | | | |
|---|---|---|---|
| Tissue infiltrate | 1 | 2 | **1.5** |
| **Overall total** | **9** | **9** | **9.0** |
| **19-054 V-nISL** | **Score** | | |
| | **Pathologist 1** | **Pathologist 2** | **Average** |
| Polymorphonuclear cells | 2 | 2 | **2.0** |
| Lymphocytes | 3 | 1 | **2.0** |
| Plasma cells | 2 | 1 | **1.5** |
| Macrophages | 3 | 3 | **3.0** |
| Giant cells | 1 | 1 | **1.0** |
| Necrosis | 1 | 1 | **1.0** |
| Capsule thickness | 3 | 4 | **3.5** |
| Tissue infiltrate | 2 | 3 | **2.5** |
| **Overall total** | **17** | **16** | **16.5** |
| **19-068 V-nISL** | **Score** | | |
| | **Pathologist 1** | **Pathologist 2** | **Average** |
| Polymorphonuclear cells | 1 | 0 | **0.5** |
| Lymphocytes | 3 | 2 | **2.5** |
| Plasma cells | 1 | 1 | **1.0** |
| Macrophages | 2 | 2 | **2.0** |
| Giant cells | 1 | 0 | **0.5** |
| Necrosis | 3 | 3 | **3.0** |
| Capsule thickness | 4 | 4 | **4.0** |
| Tissue infiltrate | 1 | 0 | **0.5** |
| **Overall total** | **16** | **12** | **14.0** |
| **19-074 V-nISL** | **Score** | | |
| | **Pathologist 1** | **Pathologist 2** | **Average** |
| Polymorphonuclear cells | 3 | 0 | **1.5** |
| Lymphocytes | 3 | 2 | **2.5** |
| Plasma cells | 1 | 0 | **0.5** |
| Macrophages | 3 | 0 | **1.5** |
| Giant cells | 1 | 0 | **0.5** |
| Necrosis | 3 | 3 | **3.0** |
| Capsule thickness | 4 | 4 | **4.0** |
| Tissue infiltrate | 2 | 3 | **2.5** |
| **Overall total** | **20** | **12** | **16.0** |

Table S3. Histopathological scores of V-nISL (PrEP) and V-PBS (control) implants. All pathologists are board-certified. Pathologist 1and 2 are from Houston Methodist Hospital and Baylor College of Medicine, respectively. V-nISL from NHP 19-087 was lost and was not scored.

| Metal | Name |
| --- | --- |
| **Dy161** | Arg1 |
| **Sm147** | aSMA |
| **Sm152** | CD163 |
| **Nd142** | CD20 |
| **Nd143** | CD3 |
| **Nd145** | CD4 |
| **Gd156** | CD45 |
| **Tb159** | CD68 |
| **Nd146** | CD8 |
| **Tm169** | Collagen |
| **Sm154** | Foxp3 |
| **Pr141** | GranzymeB |
| **Yb174** | HLA-DR |
| **Er167** | Ki-67 |
| **Dy164** | MPO |
| **Er166** | NF-kB |
| **Dy162** | NOS2 |
| **Eu151** | PD-1 |
| **Gd155** | pSTAT1 |
| **Dy163** | pSTAT3 |
| **Yb173** | t-bet |
| **Nd144** | VCAM1 |

Table S4. Markers used for IMC.
